# Mechanically primed cells transfer memory to fibrous matrices for persistent invasion

**DOI:** 10.1101/2022.05.02.490316

**Authors:** José Almeida, Jairaj Mathur, Ye Lim Lee, Bapi Sarker, Amit Pathak

## Abstract

In disease and development, cells sense and migrate across mechanically dissimilar environments. We investigated whether mechanical memory of past environments empowers cells to navigate new, three-dimensional environments. Here, we show that cells primed by stiff matrices apply higher forces, compared to soft-primed cells, to accumulate and align collagen fibers towards sustained invasion. This priming advantage persists in dense or stiffened collagen. Through an energy-minimization model, we elucidate how memory-laden cells overcome mechanosensing of softer or challenging environments via a cell-matrix transfer of memory. Consistent with model predictions, depletion of α-catenin and YAP hamper coordinated forces and cellular memory required for collagen remodeling before invasion. We release tension in collagen fibers via laser ablation and disable fiber remodeling by lysyl-oxidase inhibition; both of which disrupt cell-to-matrix transfer of memory and reduce invasion. These results have implications for cancer, fibrosis, and aging, where potential matrix memory may generate prolonged cellular response.

**One-Sentence Summary:** Cell invasion across mechanically dissimilar environments is mediated by force-based storage and extraction of cell and matrix memory.

## Introduction

To drive fundamental biological processes of development, disease, and regeneration, cells must move to and from mechanically distinct environments. For example, neural crest cells move throughout the embryo to lay the foundation for structurally complex organs (*1*) and cancer cells leave stiff tumors into softer healthy tissue to initiate metastasis (*2, 3*). Cells sense and respond to stiffness of their extracellular matrix (ECM) through actin-myosin force generation and focal adhesion signaling (*4, 5*). As a result, epithelial cells are known to migrate faster on stiffer two-dimensional (2D) surfaces (*6*). When cell collectives encounter mechanical interfaces between stiff and soft, higher cellular forces on stiffer ECM cause the whole cell colony to migrate towards the stiffer region – a process termed as durotaxis (*7*). However, it remains unknown how grouped cells negotiate such mechanical dissimilar interfaces in fibrous three-dimensional (3D) environments. If the durotaxis model shown on 2D surfaces holds true in 3D, cells moving from stiff fibrotic-like environment into softer healthy-like tissues should slow down due to a net force balance towards the stiffer side, but this has not yet been directly tested.

Further complicating this cross-environment mechanical heterogeneity, cells not only sense their current environment but also store a mechanical memory of their past environments. Indeed, stem cells have been shown to store a mechanical memory of their past stiffness on flat substrates (*5, 8*). This cellular mechanical memory is understood through a synergistic feedback between protein-level mechanotransduction, mechanosensitive transcriptional activity, and epigenetic remodeling (*9*). In the context of cell migration, as epithelial cells move from stiff to soft 2D surfaces, adequate stiff-priming enhances their migration on future soft surfaces due to a YAP-based mechanical memory (*5*). However, it is not yet clear whether such cellular mechanical memory can be imparted to collagen-rich fibrous 3D microenvironments and whether this combined cell-ECM memory can inform cell invasion (Fig. 1A).

**Figure 1.**
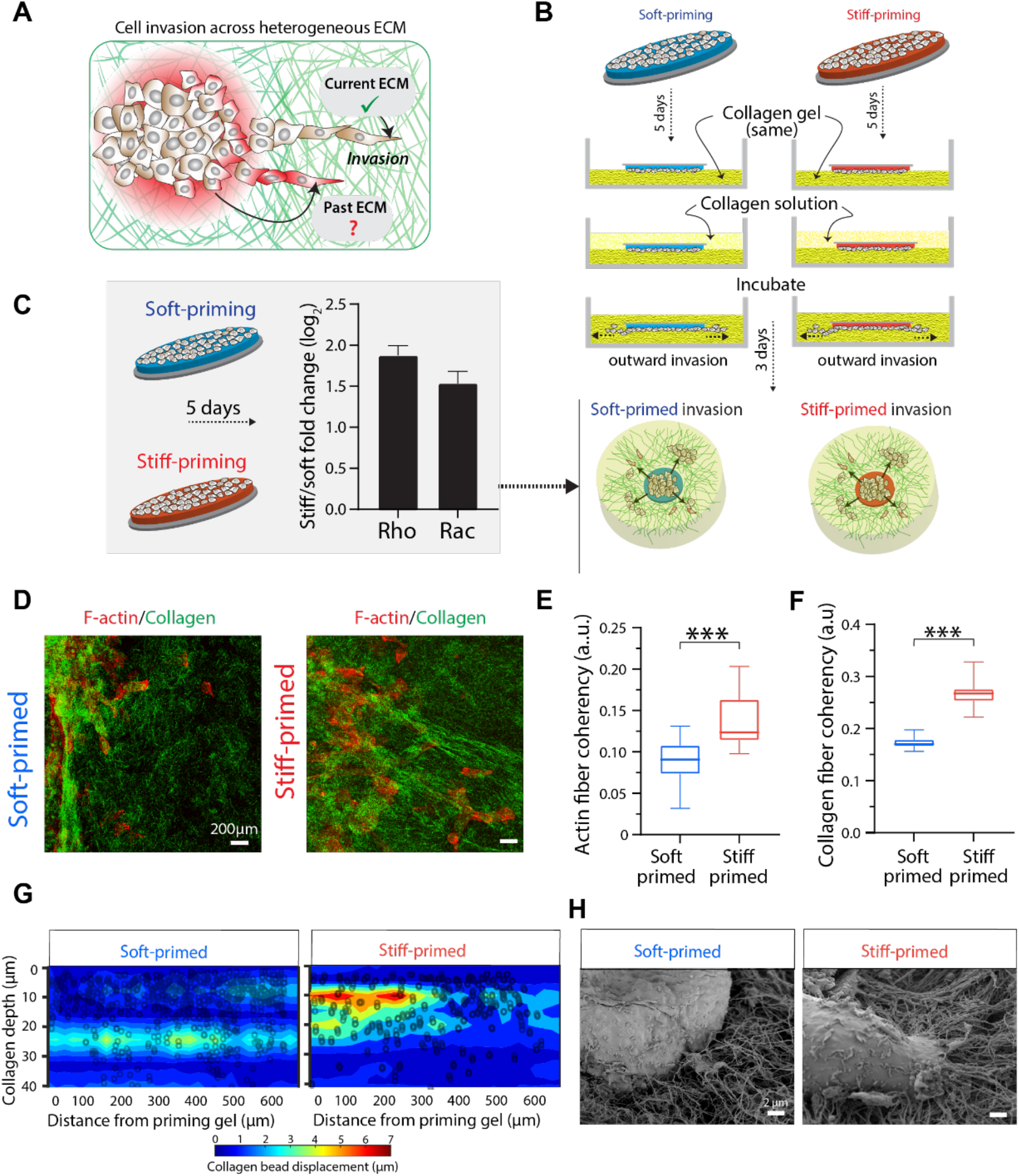
Priming of cells on stiff substrates enables higher collagen remodeling. (**A**) Schematic describing cell invasion across mechanically dissimilar environment and potential effects of current and past environments on cell invasion and matrix remodeling. (**B**) Illustration of *in vitro* device fabrication steps: cells are primed on stiff (red, 16kPa) and soft (blue, 0.08kPa) PA gels for 5 days; cell-laden PA gels are implanted into 2.3 mg/ml collagen solution; collagen gel is polymerized around primed cells; invasion of primed cells is tracked for approximately 3 days. **(C)** qPCR measurements of fold change in RhoA and Rac1 expression in stiff primed cells relative to soft primed cells after 5 days of priming. *N*=*6*. Schematic on the right describes the final dual-gel device in which differentially primed cells spontaneously invade into 3D collagen. (**D**) Representative immunofluorescence images of F-actin (red) and collagen reflectance (green) in MCF10A cells after 5 days of soft or stiff priming and 3 days of invasion into collagen (2.3 mg/ml). Scale bar, 200μm. (**E**) Coherency, a measure of alignment, of collagen and **(F)** actin fibers. *N*≥*9* RoIs. *** *P*≥*0.001*. (**G**) X-Y-Z displacements of beads in collagen matrix measured after trypsinization of cells showing higher net strain stored within collagen by stiff-primed cells. N=*5* samples. (**H**) SEM images of collagen microstructure around the invasive front of soft versus stiff primed cells. Scale bar, 2μm.

Unlike elastic hydrogels used in previous studies (*5, 9, 10*), collagen fibers undergo plastic remodeling due to cellular adhesions and forces (*11, 12*). Indeed, it has been shown that collagen deformation, alignment, and degradation are key mechanisms for cell invasion (*13, 14*). Preexisting alignment within collagen can promote cell invasion through contact guidance (*15*) and favorable collagen fiber architecture enhances cellular mechanoactivation and differentiation (*9*). Although the effect of collagen structure on cellular response is well recognized, it remains unknown whether past cellular priming can have different consequences for future collagen environments that they navigate. Since memory-laden cells remain mechanoactivated even after leaving their priming environment (*16*), it is possible that high forces of stiff-primed cells may result in greater collagen remodeling, thus transferring their memory into the matrix for follower cells to exploit for invasion. If true, cell invasion outcomes would depend not only on the current collagen structure but also on the environmental history of cells. Here, we investigate these questions through spatiotemporal invasion measurements of primed cells implanted in collagen and *in silico* modeling with an energy-minimization model for memory, mechanosensing, and collagen remodeling implemented in a lattice-based framework for cell invasion.

## Results

### Priming of cells on stiff substrates enables higher collagen remodeling

To combine cellular mechanical priming and future invasion within one assay, we developed a dual-matrix scaffold with synthetic hydrogels and 3D collagen type 1 (Fig. 1B). We first cultured MCF10A human breast epithelial cells on soft (0.08kPa) or stiff (16kPa) hydrogel substrates, which led to higher expression of known mechano-activators RhoA and Rac1 (*17*) in stiff-primed cells (Fig. 1C). Next, we implanted these hydrogel discs with adhered cells within 3D collagen (2.3 mg/ml) to allow spontaneous invasion of primed cells into a new and mechanically different environment without intermediate detachment (see Methods; Fig 1B). After 3 days of invasion through collagen (Fig. 1D; Fig S1A), we found that cells previously stiff primed had more aligned F-actin fibers (Fig. 1E; Fig S1B), compared to soft primed cells. We also measured alignment, measured as coherency, of collagen fibers around the invaded cells and found higher collagen alignment by stiff-primed cells (Fig. 1F; Fig S1C). To compare forces generated by differentially primed cells, we embedded fluorescent beads in collagen, trypsinized cells after 3 days of primed invasion, and mapped bead displacement. We found that stiff primed cells led to higher collagen deformation in the 3D space around the invasive front (Fig. 1G). Scanning Electron Microscopy (SEM) showed higher collagen bundling and accumulation around stiff-primed invading cells (Fig. 1H). Although both soft and stiff primed cells resided in the same collagen condition for ~3 days post-priming, cells that came from a stiff environment caused higher matrix remodeling in terms of both structural modifications and tension stored in collagen fibers (Fig. 1).

### Increased collagen deformation and persistent invasion by stiff-primed cells

Given the known importance cell-matrix engagement in cell migration, the differential collagen remodeling from cellular memory (Fig. 1) likely has implications for 3D cell invasion. We found that stiff priming led to almost double the invaded distance and higher number of invaded cells, compared to soft priming (Fig. 2A-C). Notably, the differences in the number of invaded cells grew larger even as cells moved farther away from the priming environment (Fig. 2C). For two other cell types, human breast cancer cell lines MDAMB231 and MCF7, we confirmed similar stiff priming advantage in 3D invasion (Fig S2). When tracked over time and distance from previous environment, stiff-primed cells pull and accumulate collagen behind the invasive front (Movie S1, Fig. 2D). These high-density collagen ‘anchors’ stay stable over the duration of observed invasion. After the first phase of ~12 hours of rapid matrix remodeling, collagen deformation rate reduced, perhaps because tension is already established within the collagen matrix (Fig. 2E). As a result, invasion speed of stiff-primed cells almost doubled in contrast to their soft-primed counterparts. Through temporal collagen bead profiles (Fig. 2D), we show that regions of high-density collagen accumulation are larger and more pronounced in case of stiff primed cells, and once formed these regions do not dissolve over time. Here, collagen deformation rate around soft-primed cells remained too low (Fig. 2E, Movie S2) to effectuate collagen accumulation or tension generation (Fig. 2D). In addition, stiff primed cells pull on collagen fibers such that deformation within the collagen matrix is spatially correlated over longer distance (Fig. S1D).

**Figure 2.**
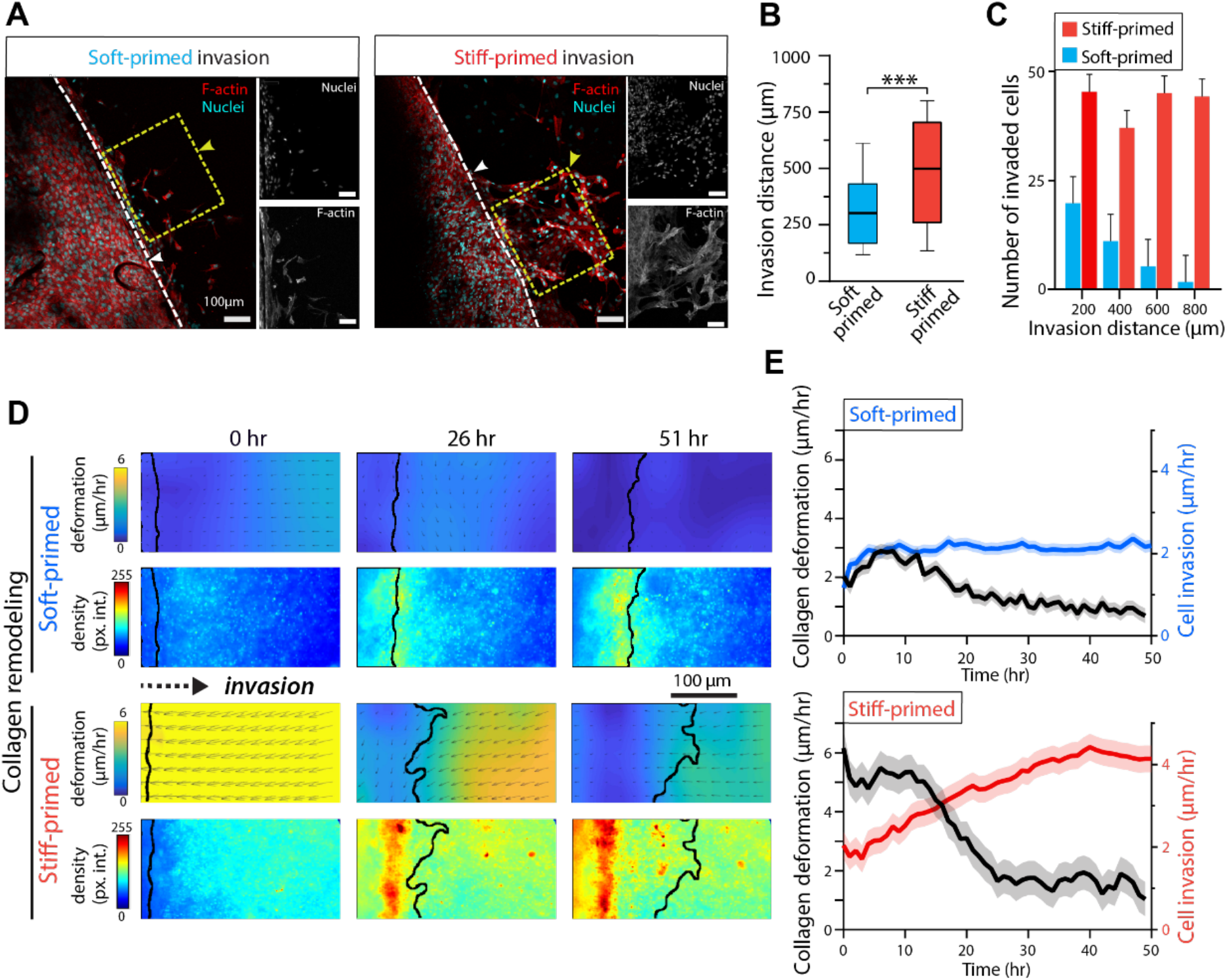
Increased collagen deformation and persistent invasion by stiff-primed cells. (**A**) At the interface between the priming PA gel and the collagen matrix, representative immunofluorescence images of MCF10A cells, F-actin (red) and nuclei (cyan), invaded after 5 days of soft or stiff priming and 3 days of invasion. White dotted lines represent the edge of the PA gel. Yellow dotted line represents RoIs of cell invasion. Scale bar, 100μm. (**B**) Average invasion distance, *N*=*5*, \*\*\**P*≤*0.001*, and (**C**) average number of cells relative to increasing distance of invasion after stiff (red) and soft (blue) priming. *N*=*5*. (**D**) Heatmaps of collagen deformation rate (PIV vectors) and collagen bead intensity at three selected timepoints, showing collagen remodeling by soft or stiff primed cells (black outline annotates the cell invasion front). The middle timepoint (*t*=*26 hr*) indicates that majority of collagen accumulation occurs in the first half of the net invasion period while major cell invasion follows in the second half. Dotted arrows show direction of invasion. Scale bar, 100μm. (**E**) Rates (mean ± SEM) of collagen deformation and cell invasion over time caused by soft or stiff primed cells; stiff-primed cell invasion increases after collagen deformation reduces. *N*=*8*.

Given the large-scale collagen remodeling (spanning hundreds of microns) over days and the clear role of cell-generated forces, we next asked whether such forces need to be collectively generated by the cell population. Upon depleting α-catenin, a cell-cell adhesion protein, in MCF10A cells, the resulting collagen deformation and cell invasion rates became uncoordinated and priming-independent (Fig. S4).

### Differential kinetics of cellular memory, mechanosensing, and matrix remodeling explains priming-dependent cell invasion

Thus far, experiments have revealed that collective cellular forces of past stiff priming can install the cellular mechanical memory into the matrix through plastic collagen remodeling, defined by accumulation, alignment, and tension, which helps propel future cell invasion. As cells enter a 3D matrix, resistance (*γ*) due to matrix density must be overcome via collagen remodeling (*α*) to enable invasion. We define the net mechano-activated migration potential from effective protrusions, –Δ*H_protrusions_* = *μ* as the sum of current cell mechanoactivation (*ϕ*), collagen remodeling (*α*), and overall matrix resistance (*γ*) (Fig. 3A). We model that collagen remodeling occurs due to two cell signals: (1) acquired mechanical memory (*ψ*) by the past environment and (2) direct mechanosensing (*ϕ*) of the current matrix (Fig. 3B). As cells face bulk resistance from the newly encountered collagen matrix, collagen remodeling serves as a ‘transfer’ of mechanical memory from cells to the ECM, which cells then utilize to invade. According to our model, mechanical memory decays slowly, consistent with previous studies (*9, 17*), and direct mechanosensing (*ϕ*) adapts quickly.

**Figure 3.**
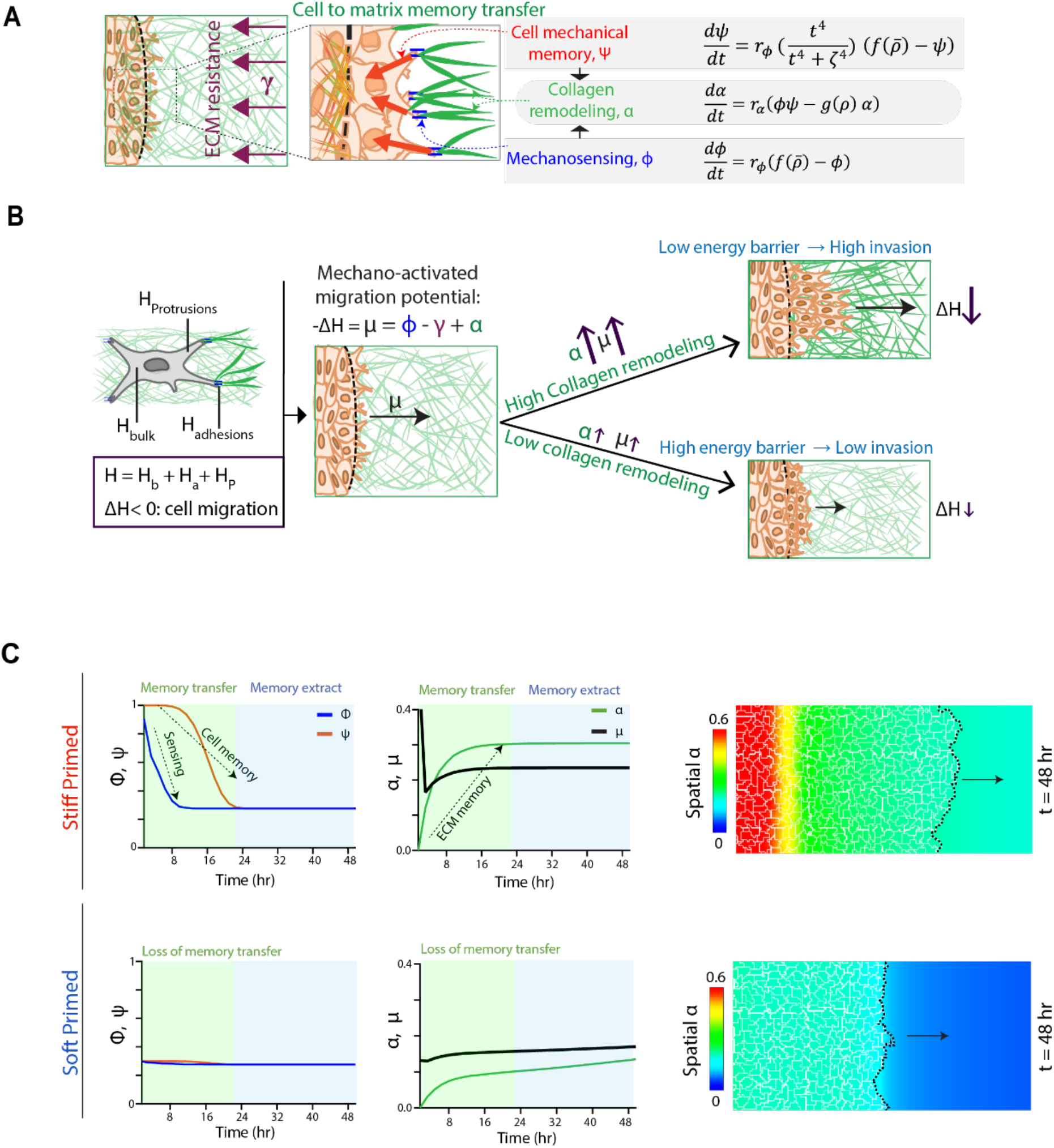
Differential kinetics of cellular memory, mechanosensing, and matrix remodeling explains priming-dependent cell invasion. **(A)** Modeling scheme for primed cell invasion into 3D collagen matrix – as memory-laden cells move into 3D matrix, the invasion is opposed by ECM resistance *γ*, mechanical memory (*ψ*) and direct mechanosensing (*ϕ*) of cells perform collagen remodeling *α*, which lower the energy required for net protrusions and invasion μ. **(B)** As a cell migrates, it has to extend protrusions, form adhesions and break rear adhesions, all of which have an energetic cost associated. The energies associated with adhesions (*H_adhesions_*), bulk cell properties (*H_cytoskeleton_*) and protrusions (*H_protrusions_*) are shown here. To minimize its net energy, the cell would only move forward when net change in energy Δ*H* < 0. We define the mechano-activated net protrusions as –Δ*H_protrusions_* = *μ* as the sum of current mechanoactivation (*ϕ*), remodeling (*α*) and resistance (*γ*). Collagen remodeling causes an increase in net protrusive potential, which lowers Δ*H*, making invasion more favorable. Low collagen remodeling keeps the barrier high and will cause less invasion **(C)** Temporal evolution of key signals as primed cells invade through collagen: direct mechanosensing of current collagen matrix *ϕ* and cellular memory *ψ* (left column); net collagen remodeling 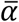 and net protrusions *μ* (middle column); spatial heatmap of remodeled collagen *α* and state of the invasive front (black dotted line) at the final timepoint of *t* = *48 hr* (right column).

To better understand how cell and matrix memories work in conjunction with conventional mechanosensitivity, we start with classic models of cell movement resulting from protrusion, contraction, and adhesive resistance (*18*) (Fig. 3B). As a cell migrates, it has to extend protrusions, form new adhesions, and break rear adhesions, all of which have an associated energetic cost. Here, we define total energy of the system (*H*) as the sum of the energies associated with adhesions (*H_adhesions_*), bulk cell properties (*H_bulk_*) and protrusions (*H_protrusions_*) (Fig. 3B). We model the cell to minimize its energy, and it would only move forward when the net change in energy Δ*H* < 0 (Fig. 3B). Collagen remodeling causes an increase in the net migration potential, which lowers Δ*H*, making invasion more favorable. Low collagen remodeling keeps the energy barrier high and will cause less invasion (Fig. 3B). We implemented this energy balance of mechanosensing (*ϕ*), resistance (*γ*), and memory-driven collagen remodeling (*α*) resulting in a net migration potential *μ* in a lattice-based model for cell invasion (*SI Appendix*).

To explain and understand experimental results, we simulated spatial and temporal evolution of cell colonies that were stiff-primed (*ϕ* = *ψ* = 1) and soft-primed (*ϕ* = *ψ* = 0.3), at *t* = 0, entering collagen matrix (*ρ* = 2.3 *mg/ml*). As cells enter 3D collagen, the mechanosensing signal (*ϕ*) quickly adapts to the current soft signal received from the new collagen environment, and the net migration potential (*μ*) drops due to the ECM resistance (*γ*) (Fig. 3C, top row). However, cellular mechanical memory (*ψ*) stays high for ~12 hours, allowing collagen remodeling *α* to rise (Fig. 3C, top row) and growing the size of collagen ‘anchors’ (defined by high intensity regions of *α*; Fig. S3A). Collagen remodeling (*α*) reduces the resistance to invasion and causes a rise in net migration potential (*μ*). After 12 hours, although the cellular memory *ψ* adapts to the new soft collagen, the remodeled matrix (*α*) continues to work against the ECM resistance to maintain high net migration potential *μ*, which results in cell invasion going forward (Fig. 3C, top row; Movie S3). In contrast, soft primed cells do not carry such cellular mechanical memory that can be used for collagen remodeling, and thus they cannot overcome the ECM resistance. As a result, net migration potential (*μ*) does not rise, leading to low invasion (Fig. 3C, bottom row; Movie S3). Thus, the transfer of memory from cells to the matrix is a crucial step in the cross-environment collective invasion of epithelial cells into collagen.

### Loss of cellular memory obviates matrix memory and priming-dependent invasion

Overall, this model merges the complex interplay between memory kinetics, direct mechanosensing of immediate matrix, and collagen remodeling with the conventional models of mechanosensitive cell migration (*18, 19*). Given the importance of priming in maintaining cellular memory, we asked whether stiff primed cells would continue to invade if cells could not sustain mechanical memory. In our model, the rate of decay of cellular memory is captured by parameter *ζ* whose higher values ensure more stable memory (Fig. 4A). We simulated cells primed for 5 days on stiff substrates followed by invasion into collagen, but their cellular memory was allowed to deplete rapidly (*ζ* = 50) (Fig. 4A-C). Our simulations show that rapid depletion of cellular memory (*ψ*) does not allow enough collagen remodeling (*α*) required to overcome ECM resistance (Fig. 4B,C). As a result, memory-depleted cells invaded collagen at a slower speed compared to those with stable memory, despite prior stiff priming in both cases (Fig. 4B,C).

**Figure 4.**
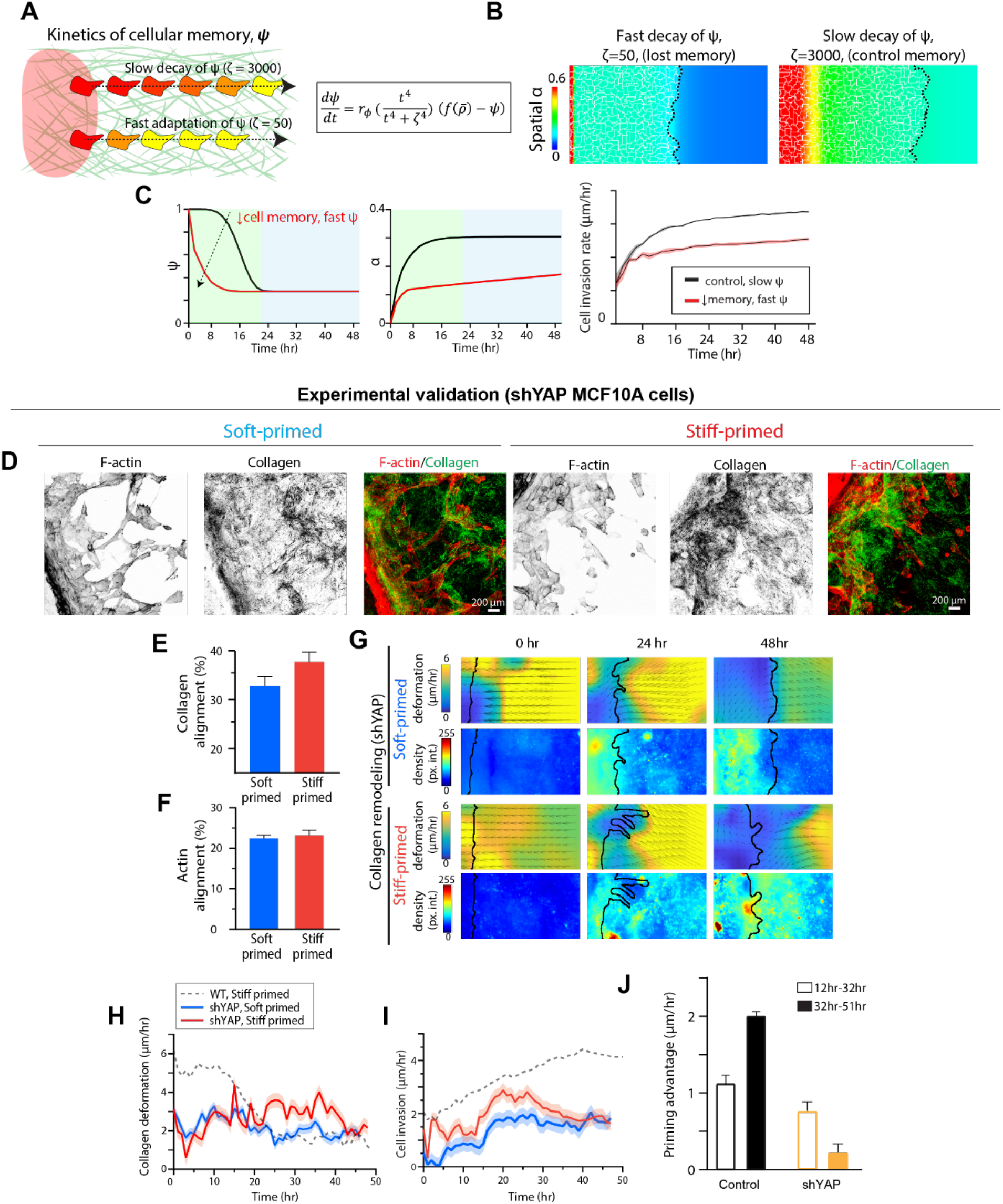
Loss of cellular memory obviates matrix memory and priming-dependent invasion. **(A)** Schematic describing the role of cellular memory kinetics by varying the parameter ζ whose lower values expedite memory decay. **(B)** Spatial distribution of collagen remodeling and invasion of stiff primed cells for two cases: depleted of cellular memory (left) and the control case (right). **(C)** With an expedited decay rate, simulations predict that ψ adapts quickly to the current collagen (red line), causing reduction in collagen remodeling α, and slowed cell invasion rate. **(D)** Split channel images of collagen reflectance and F-actin immunofluorescence along with merged image for shYAP primed cell invasion. Scale bar, 200μm. **(E)** Average percentage of collagen fiber alignment caused by stiff (red) and soft (blue) primed shYAP cells. *N*=*8*. **(F)** Average percentage of actin fiber alignment on both stiff (red) and soft (blue) primed shYAP cells. *N*=*8*. **(G)** Heatmaps of collagen deformation (PIV vectors) and collagen bead intensity caused by soft or stiff primed shYAP cells (black outline annotates the cell invasion front). **(H)** Rates of collagen deformation and **(I)** cell invasion over time caused by soft or stiff primed shYAP cells. Here, the dotted gray line represents corresponding data for stiff primed wild type cells. *N*=*4*. **(J)** Average priming advantage (difference between cells invasion rates of stiff and soft primed cells) of shYAP cells.

Thus, two different kinetics in cellular mechano-sensation signaling – slow decay of mechanical memory and fast adaptation to current environment – are both required for seamless transfer of memory from cells to matrices as well as direct mechanosensing of current ECM for priming-dependent invasion. To test our model of matrix remodeling coupled with cellular memory, we depleted yes-associated protein (YAP) – a known memory regulator (*9, 20*) – in MCF10A cells and repeated experiments of primed cell invasion. Here, MCF10A-shYAP cells were stiff or soft primed, and allowed to invade 3D collagen. We found that there was no substantial difference between soft and stiff primed cells in terms of alignment of F-actin fibers and collagen fibers (Fig. 4D-F). As cells invade, the resulting collagen deformation was priming-independent, somewhat temporally uncoordinated, and lower compared to the control (Fig. 4G,H). Consistent with the model that early collagen remodeling is required for future invasion, lower collagen deformation of shYAP cells coincided with slower invasion speeds (Fig. 4I), regardless of soft or stiff priming. To quantify the effect of past priming on future cell invasion, we calculated a ‘priming advantage’ defined by the difference in the invasion speeds of soft and stiff primed cells. While this priming advantage increased over time for control cells, it was significantly lowered in case of shYAP (Fig. 4J). Overall, our model and the experimental validation with shYAP show that the loss of stable cellular memory eliminates the possibility of a matrix memory, resulting in priming-independent cell invasion.

### Priming advantage towards cell invasion persists in dense or stiffened collagen

Since dense or crosslinked collagen matrices are known to restrict cell invasion (*12, 21*), we asked whether mechanical memory could prove advantageous to cells when they encounter such challenging environments (Fig. 5A). We repeated experiments in two collagen compositions: first of higher collagen density of 3.1 mg/ml, and second with increased stiffness but constant density by mixing 2.3 mg/ml concentration of collagen (same as control) mixed with Riboflavin, a known photo-crosslinker, followed by UV exposure to photo-activate crosslinking of collagen (*22, 23*). We measured stiffness of collagen gels using Atomic Force Microscopy (AFM) and found that increasing collagen density to 3.1 mg/ml almost doubled the stiffness to an average of ~1kPa, compared to ~0.5kPa stiffness for control collagen density of 2.3 mg/ml (Fig. 5B). Photo-crosslinked collagen led to an even higher increase to ~2.4kPa average stiffness (Fig. 5B). To understand how differentially primed cells behave in these collagen matrices, we allowed soft and stiff primed cells to invade, imaged collagen fibers (Fig. 5C), and found higher cell invasion and fiber alignment (Fig. 5D, Fig S5A-B) by stiff primed cells. Compared to the control collagen concentration, higher density and crosslinked collagen matrices led to overall slower invasion and lower collagen deformation (Figs. 5E,F left column). Even so, the stiff-primed cells generated higher collagen deformation rate (Fig. 5E right column, Fig. S6) and resulted in faster cell invasion compared to soft priming (Fig. 5F, right column). Although cell invasion rate reduced in these challenging matrices of higher density and crosslinking, the priming advantage towards invasion persisted (Fig. 5G). Through collagen bead intensity kymographs (Fig. 5H, Fig. S6), we show that stiff primed cell generated higher collagen accumulation in both dense and crosslinked collagen. To test whether our computational model can capture these experimental findings of priming-dependent invasion in varying collagen compositions, we repeated simulations with higher collagen density (*ρ* = 3.1 *mg/ml*) and crosslinked collagen (*ρ* = 2.3 *mg/ml*, 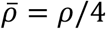, *γ*_0_ = 0.06). These parameters represent the collagen configuration for riboflavin treatment, which increases collagen stiffness (Fig. 5B), makes fibers thicker (Fig. S5C), and reduces the pore size (Fig. S5D). Consistent with experiments, although collagen deformation and invasion rates were reduced in dense and crosslinked collagen, stiff-primed cells continued to perform better than soft primed cells (Fig. 5I). Overall, our experiments and simulations show that prior mechanical priming continues to be advantageous in denser and stiffened collagen matrices due to memory-dependent collagen remodeling.

**Figure 5.**
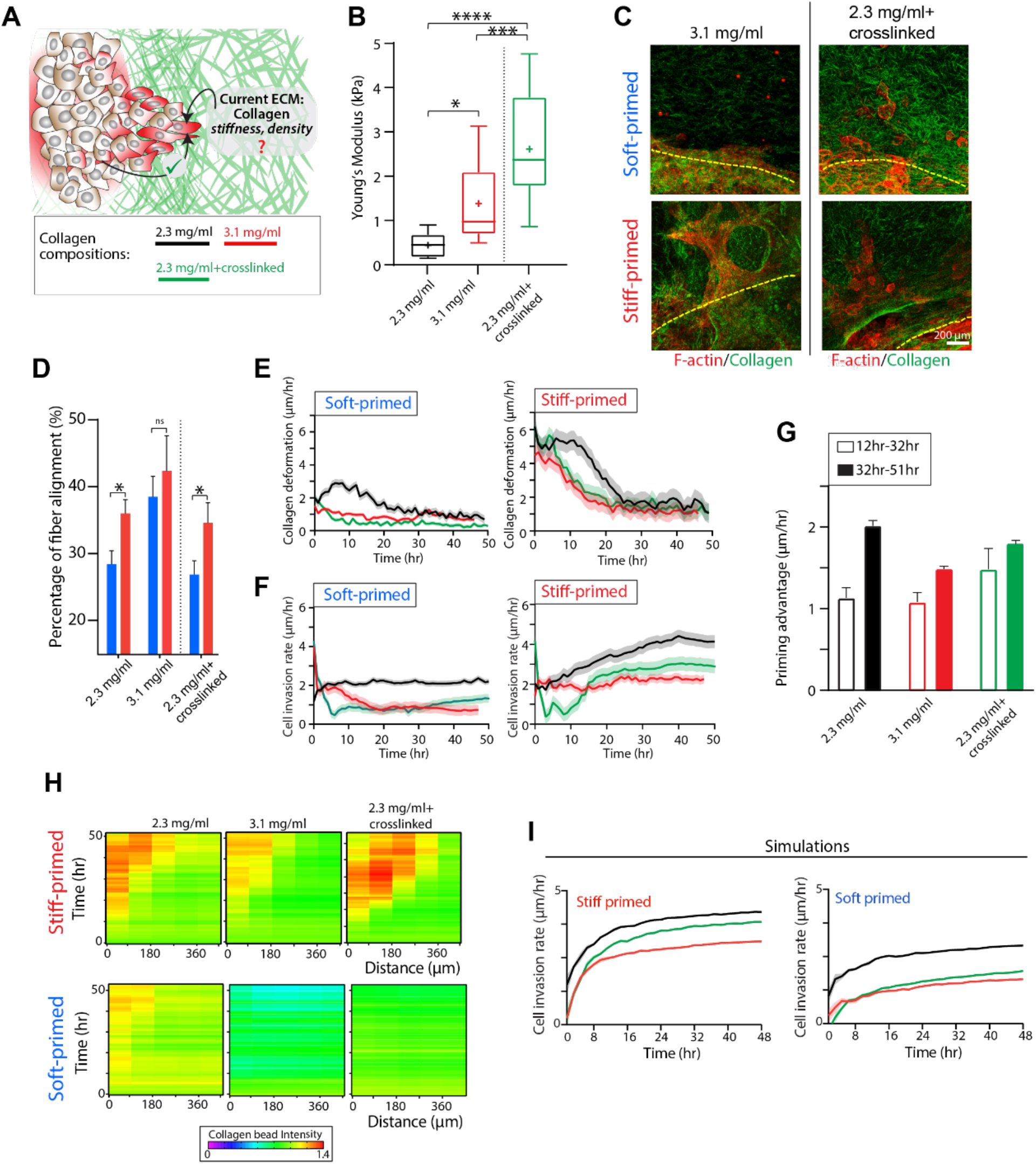
Priming advantage towards cell invasion persists in dense or stiffened collagen. **(A)** Schematic posing a question of how differentially primed cells navigate different collagen compositions defined by density and crosslinking, 2.3 mg/ml, 3.1 mg/ml, and 2.3 mg/ml with crosslinking. **(B)** Average Young’s Modulus measured from AFM measurements of collagen gels from three different formulations: 2.3 mg/ml (control), 3.1 mg/ml (dense), and 2.3mg/ml + crosslinked using Riboflavin (crosslinked). *N*=14. *** *P*≤*0.001*, * *P*≤*0.03*. **(C)** Immunofluorescence images of MCF10A cells invaded into 3.1 mg/ml collagen and 2.3 mg/ml+crosslinked collagen after 5 days of soft or stiff priming followed by 3 days of invasion; F-actin (red), collagen (green). Yellow dotted line shows the edge of the PA gel. Scale bar, 200μm. **(D)** Average percentage of fiber alignment for both stiff primed (red) and soft primed (blue) cells invading through control, dense, and crosslinked matrices. *N*= *6*. * *P*≤0.03, *ns* = *not significant*. **(E)** Rate of collagen deformation and **(F)** cell invasion over time caused by soft or stiff primed in control, dense, and crosslinked collagen *N* = *8*. **(G)** Average priming advantage (difference between invasion rate of stiff- and soft-primed cells) for control, dense collagen, and crosslinked collagen conditions. Comparison between the first phases of invasion (outlined box; 12-32 hr), which is dominated by collagen remodeling (Fig. 2), and the second phase of invasion (filled box; 32-51 hr) shows that the “priming advantage” increases or stays steady over time across collagen conditions. *N* = *8*. **(H)** Kymographs of collagen bead intensity over time of invasion and distance from the PA gel for soft and stiff primed cells in control, dense, and crosslinked collagen. *N*=*8*. Here, the red band of high bead intensity in case of stiff primed cells indicates collagen accumulation that is sustained over the duration of invasion analysis (50 hr). **(I)** Cell invasion rates for stiff primed and soft primed cells calculated from simulations for control (black line), dense collagen (red line) and stiffened collagen (green line)

### Disruption of stable collagen remodeling abrogates the effect of past cellular priming on future invasion

According to our model and results thus far, memory-dependent cellular forces pull on collagen fibers to enable cell invasion. This collagen remodeling needs to be stable over time such that cells can continue to move as they go from one environment to another (Fig 6A). To test the effect of collagen remodeling kinetics on invasion, we first used our model to vary rate constant *r_α_* that determines the speed of collagen remodeling (Fig. 6B) and its smaller values led to lower levels of remodeled collagen (*α*) by stiff-primed cells (Fig. 6C). If the matrix remodeling is not performed fast enough, before the cellular memory dissipates, the matrix memory is not stored. Since remodeled collagen is required to reduce the energy barrier for cell invasion, lower levels of collagen remodeling (due to lower values of *r_α_*) led to slower cell invasion (Fig. 6C).

**Figure 6.**
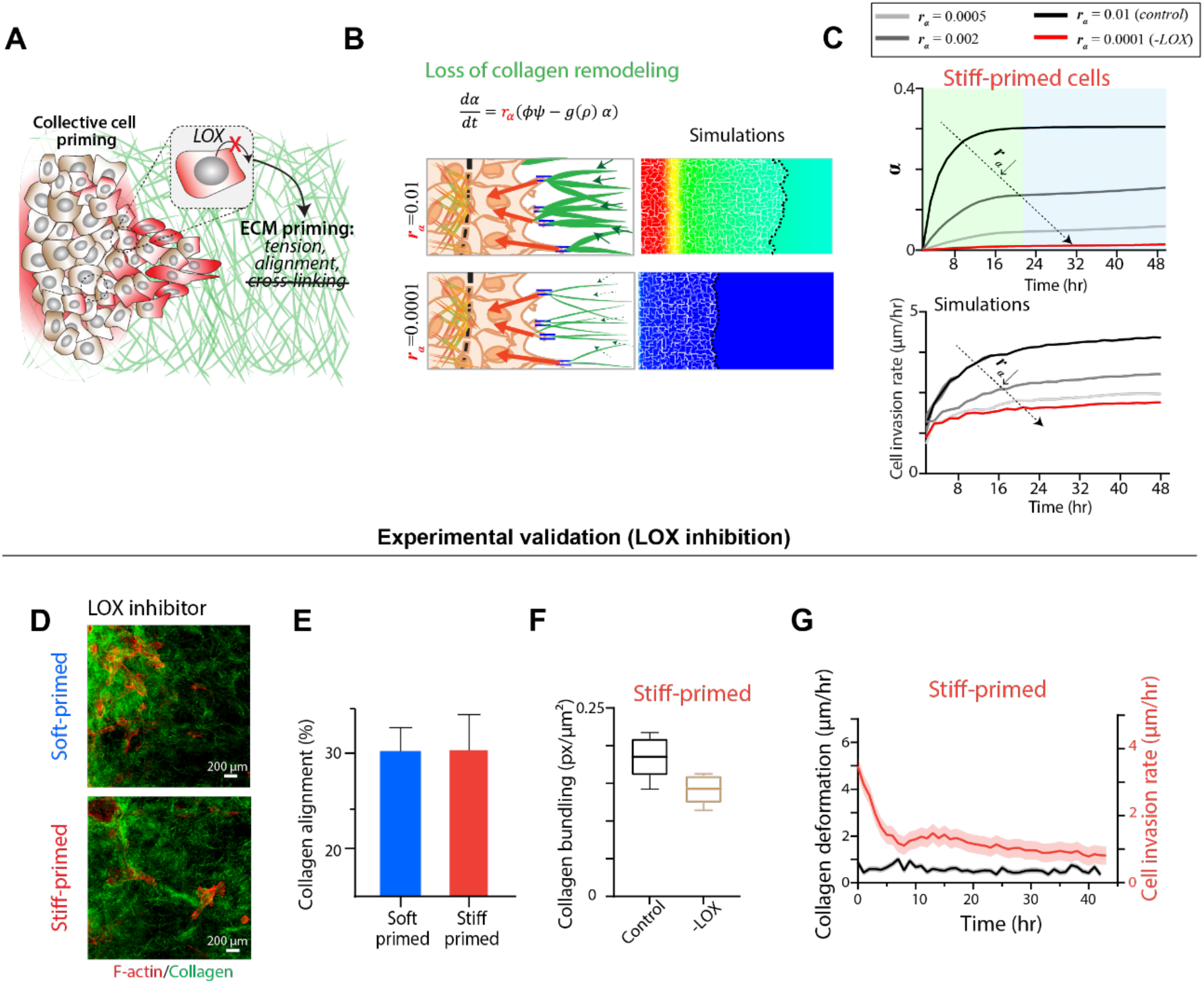
Disruption of stable collagen remodeling abrogates the effect of priming on invasion. **(A)** Schematic posing the question of how the loss of collagen crosslinking could disrupt priming-dependent invasion. **(B)** Schematic and simulation results showing spatial map of remodeled collagen, α, and cell invasion at final timepoint (*t* = 48 *hr*) for two different rate constants (*r_α_*) for collagen remodeling kinetics. **(C)** Simulated net collagen remodeling and cell invasion rate over time for varying values of *r_α_* = (0.01,0.002,0.0005,0.0001 *h*^−1^). **(D)** Representative immunofluorescence images of stiff- and soft-primed cells treated with BAPN, LOX inhibitor, showing F-actin (red) and collagen reflectance (green) after primed invasion. Scale bar, 200μm. **(E)** Percentage of aligned collagen fibers, *N*=*4*, in soft- and stiff-primed LOX-inhibited cells. **(F)** Collagen bundling and **(G)** temporal rates of collagen deformation and cell invasion caused by stiff-primed cells with and without LOX inhibitor. *N*=*4*.

To experimentally test the role of stable collagen remodeling in primed invasion, we used β-aminopropionitrile (BAPN) to inhibit lysyl oxidase (LOX) that is known to crosslink collagen (*24*). We expect that the loss of crosslinking would slow collagen remodeling. After soft or stiff priming of cells, BAPN was added, and cells were allowed to invade into collagen (Fig. 6D; Fig. S7A). Through collagen fiber imaging and analysis, we found that collagen fiber alignment was independent of past soft or stiff priming (Fig. 6E, Fig. S7B). In case of stiff primed cells, LOX inhibition reduced collagen fiber bundling compared to the control, untreated case (Fig. 6F). These structural analyses of collagen fibers (Figs. 6E,F) indicate that LOX inhibition reduces collagen remodeling after differential priming of cells (Fig. S7C). When tracked over time (~2 days), collagen deformation remained low and invasion rate of stiff-primed cells remained low (Fig. 6G), which is consistent with modeling predictions.

### Spatial propagation of collagen remodeling is required for priming-dependent invasion

The accumulation of collagen remodeling depends on collective and spatially coordinated cellular forces (Figs. 1,2). As we have shown (Fig. 4; Fig. S4), the ability of cells to generate forces and retain them via mechanical memory are important for invasion. We next asked whether cellular forces are just necessary around the cells, or they need to be propagated through the matrix to enable collagen remodeling (Fig. 7A). In our model, after primed cells reach collagen, they not only remodel their surrounding collagen, but the energy field must spatially propagate through the collagen matrix, capturing force propagation, to enable sustained cell invasion that we see in experiments. To test the effect of spatial collagen remodeling, we performed simulations of primed cell invasion with reduced distance of propagation of collagen remodeling *α* (parameter *n* = 1; Fig. 7B). Higher value of *n* caused a reduction in net accumulation of remodeled collagen (*α*). In this case, since invasive front of stiff-primed cells is unable to cause large-scale changes in the matrix and thus store their memory, the net cell invasion rate reduced (Fig. 7C). To implement the similar spatial disruption of force-based collagen remodeling in experiments, after the stiff-primed cells invaded into the collagen we performed laser ablation of collagen fibers ahead of the invasive front, at the onset of invasion. As shown in a represented image (Fig. 7D), collagen fibers in contact with the invading cells are intact; however, they are disconnected from the rest of the collagen matrix. In this case, overall collagen deformation was substantially reduced within a few hours after laser ablation of fibers and remained in a low net deformation state over time (Fig. 7E; Fig. S8), which indicates the importance of long-distance force propagation within collagen for bulk remodeling. Similarly, although the invasion rate of stiff-primed cells started at a high level, it subsided within 6-8 hours and remained low over two days of tracking (Fig. 7F). In sum, just as the stable memory and collective forces are important in cells (Fig. 4, Fig. S4), stable remodeling and long-distance force propagation are important in collagen matrices for the stated cell-matrix memory transfer and priming-dependent cell invasion.

**Figure 7.**
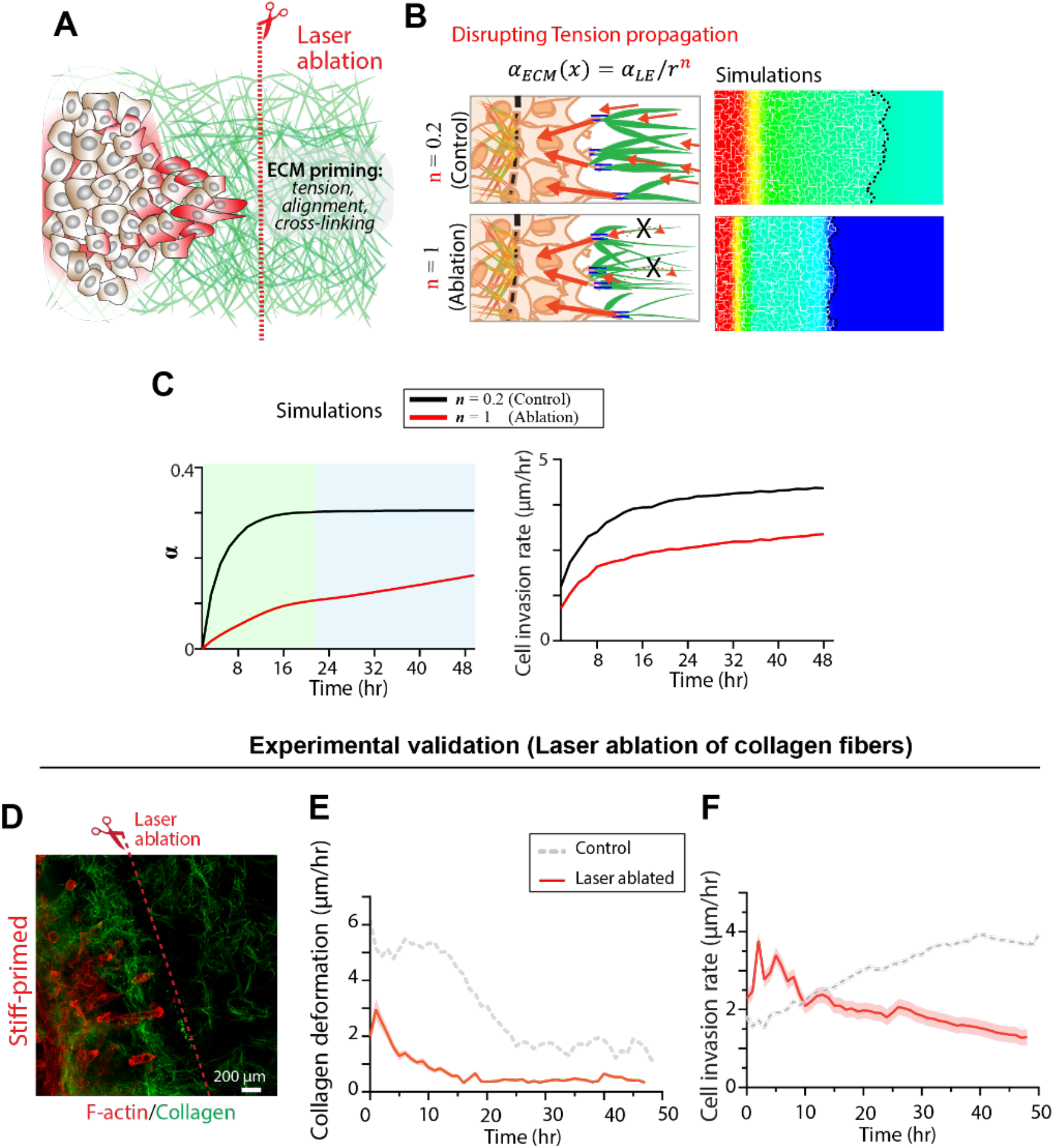
Spatial propagation collagen remodeling is required for priming-dependent invasion. **(A)** Schematic of laser ablation of collagen fibers ahead of the invasive front. **(B)** Effect of reduced distance of tension propagation within collagen, for two different cases, shown schematically and through simulated spatial map of remodeled collagen, *α*, and cell invasion at final timepoint (*t* = 48 *hr*). **(C)** Simulated net collagen remodeling and cell invasion rate over time for comparing the control case against the case with ablated propagation of remodeling across collagen. **(D)** Representative immunofluorescence image showing laser ablation of collagen fibers ahead of the invasive front of stiff-primed cells, with F-actin (red) and collagen reflectance (green). Scale bar, 200μm. **(E)** Temporal rates of collagen deformation and **(F)** cell invasion, caused by stiff-primed cells with and without laser ablation of collagen fibers. *N*=*4*.

## Discussion

Collagen remodeling by cell-generated forces and reinforcement of cellular mechano-response by matrix mechanics have been known and studied for several decades. Upregulation of actin-myosin contractility and Rho signaling allow the cells to either squeeze through narrow pores or align collagen fibers to make paths in three-dimensional environments (*25, 26*). In recent years, cells’ ability to store mechanical memory of their past environments has emerged, which can sustain cellular forces even after they arrive in soft environments, as we have previously shown on 2D surfaces (*9*). Here, we asked whether memory-derived cellular forces can alter future matrices, which is only possible in case of fibrous 3D matrices not 2D linear elastic substrates. As a result, mechanical memory could remodel the matrix state and thus alter the course of future invasion, as cells move across mechanically distinct environments. We developed a novel hydrogel-collagen device and analysis procedure to combine mechanical memory and cross-environment invasion within one system. Here, we show that cells previously primed by stiff environments continue to invade through softer 3D collagen matrices, mediated by fiber remodeling.

Given previous studies on cellular mechanical memory (*9, 20*), it is not surprising that past priming on stiff substrates enhances future cell migration; rather, the new insights are attributed to the discovery that memory-dependent invasion is temporally and spatially coupled with irreversible collagen remodeling generated by collective cellular forces. By mapping collagen deformation and cell invasion over time and space, we showed that much of the collagen remodeling happens before the outward cell invasion (Fig. 2). During this process, cells accumulate collagen ‘anchors’ behind the invasive front and then use the aligned and stressed collagen environment to further invade. As such, collagen remodeling and cell invasion are temporally staggered processes – high cellular forces carried over by mechanical memory are exploited first in remodeling the matrix, which we refer to as a ‘transfer of matrix memory’, and then cells invade by utilizing this remodeled matrix. This bulk-scale matrix remodeling depends on collective cellular forces, as evident by the loss of collagen deformation after cell-cell adhesions are depleted via α-catenin. To test the possibility that our observations are due to some mechanical artifacts of an interface between stiff hydrogel and soft collagen, we used memory-null YAP-depleted cells, which do not undergo priming-dependent collagen remodeling or invasion despite crossing the same interface as the wildtype cells.

Our results show that fibrous microenvironment does not just act as a passive provider of mechanical cues; instead, memory-laden cells can actively remodel the matrix for persistent invasion into new environments. These findings add to the existing concepts of cellular mechanosensing via direct adhesions with the immediate matrix. In cell migration on flat surfaces with a stiffness gradient, the process of durotaxis allows preferential cell migration towards stiffer regions, due to intra- and inter-cellular force propagation (*7, 27*), which also occurs *in vivo* along cell-generated stiffness gradients (*28*). According to our results, when the new environment is fibrous and plastic, albeit soft, cells are not bound by the durotaxis model, because the matrix not only governs cellular mechanosensing but also stores cells’ memory, causing persistent cell invasion despite the preexisting stiff-to-soft gradient. We explain this process through mechanisms analogous to the known rudimentary steps in classic neurological memory – encoding, storage, and retrieval (*28*). To better understand the temporal and spatial coupling of cell and matrix memory, we developed a mathematical model in three key steps of memory-dependent invasion: encoding prior mechanical priming of cells in the form of sustained mechanotransduction, e.g., YAP (*9, 20*); storage of memory into the matrix via force-based fiber remodeling; and retrieval or extraction of matrix memory by cells invading through the remodeled collagen. Our model combines individual influences of direct mechanosensing and acquired memory through their differential kinetics. Through simulations and experimental validation, we show that memory-based cellular forces and matrix remodeling are such delicately coupled processes that any disruptions in collagen crosslinking or force propagation remove the effect of past cellular priming on future invasion. That being said, without extraneous disruptions to the system, the fundamental mechanisms of cell-to-matrix transfer of memory and invasion hold true for multiple cell types and collagen compositions. Overall, our results show that the past ‘stiff-priming’ provides cells a previously unappreciated advantage towards navigating new challenging environments.

Although stem cell mechanical memory and cellular reprogramming had previously been shown (*5, 20, 29*), our findings reveal that such memory can be encoded into the ECM. In stem cells, cardiac cells and breast cancer cells, mechanical priming can result in chromatin remodeling (*30–32*), causing epigenetic modifications and potentially mechanical memory. Given that our model captures fundamental mechanisms of coupling between cellular and matrix memory using their differential kinetics within an energy-minimization framework, it can be adapted in future studies to develop unifying principles of mechanical memory across scales. We also speculate that while the biochemically stored cellular memory may deplete due to changing transcriptional and epigenetic landscapes, the matrix memory is mechanical in nature and thus may have long-lasting effects on cellular response. Our experimental and mathematical models combining cellular and ECM memory broadens the previously understood processes of collective cell invasion in complex fibrous environments. This work highlights the need to account for both past and current cellular state and ECM remodeling in heterogenous cell invasion. Our findings of cell-to-matrix transfer of mechanical memory adds a new consideration in wide-ranging biological processes wherever cells change environments, e.g., fibrosis, cancer, and aging.

## Materials and Methods

### Cell culture and reagents

MCF10A (ATCC), non-tumorigenic human breast epithelial cells, and its variants anti-YAP shRNA (shYAP) and anti–α-catenin shRNA (α-cat-KD) (both courtesy of G.D. Longmore, Washington University) were cultured in DMEM/F12 (GE Healthcare Life Sciences), supplemented with 5 % horse serum (Invitrogen), 20 ng/ml epidermal growth factor (EGF, Miltenyi Biotec Inc), 0.5 mg/ml hydrocortisone (Sigma-Aldrich), 10 μg/ml insulin (Sigma-Aldrich), 100 ng/ml cholera toxin (Sigma-Aldrich) and 0.2 % Normocin (Invitrogen). Media was changed every 3 days while cells were in culture plate.

### shRNA knockdown

shRNA knockdowns of YAP and α-catenin were chosen to disrupt stiffness sensitivity and cell-cell communication, respectively. A healthy MCF10A cell line was depleted of YAP using lentiviral pFLRU vector containing anti-YAP shRNA, developed and verified previously by us (*9*). Depletion of α-catenin in healthy MCF10A was done by lentiviral pFLRu vector containing anti–α-catenin shRNA, as previously used (*33*).

### Polyacrylamide gel preparation

Glass coverslips of 5 mm diameter (Thermo Fisher Scientific) and glass slides were prepared for gel adhesion. Coverslips were activated by plasma cleaning and 10 min treatment with bind-silane solution, composed of 94.7% ethanol, 5% acetic acid, and 0.3% bind-silane (GE Healthcare Life Sciences). Following treatment, coverslips were washed with ethanol and air-dried. While waiting for the coverslips to dry, glass slides were treated with Sigmacoat solution (Millipore) to create a hydrophobic surface. Polyacrylamide gels with distinct stiffness were fabricated through step-by-step polymerization of PA solution. Briefly, precursor solutions combining acrylamide (A; Bio-Rad), bis-acrylamide (B; Bio-Rad), and ultrapure water were mixed for respective contents of 3%, 0.05%, and 96.95% (~0.08 kPa stiffness) and 12%, 0.15%, and 87.85% (~16 kPa stiffness). These solutions were degassed by N2 injection. Then, volumes of ammonium persulphate (APS) and N, N, N′, N′-tetramethylethylene diamine (TEMED) were added to yield concentrations in the final gel solution of 0.5 and 0.05 %, respectively. To form gels, this solution was sandwiched between the treated glass slides and 5 mm coverslips. These constructs were placed in a degasser for 15 min to polymerize, then were submerged in DPBS (Gibco) for 30 min. Coverslips were lifted, and formed gels were placed for 1 hour under UV for sterilization. To facilitate collagen conjugation, gels were functionalized with 0.5 mg/ml solution of sulfosuccinimidyl 6-(4′-azido-2′-nitrophenylamino) hexanoate (Sulfo-SANPAH) (Thermo Fisher Scientific) prepared in 50 mM HEPES buffer (Santa Cruz Biotechnologies). To adhere Sulfo-SANPAH to PA gel surfaces and provide binding sites for collagen, Sulfo-SANPAH coated gels were exposed to 365 nm UV for 10 min. Gels were then washed twice before coating with 0.05 mg/ml collagen type I (rat tail, Santa Cruz Biotechnologies). Finally, to achieve sufficient collagen adhesion, gels were incubated in this solution overnight at 4°C.

### Mechanical priming of cells and collagen gel fabrication

To mechanically prime cell colonies before their invasion, cells were plated on PA gels of different stiffnesses (~0.08 kPa and ~16 kPa). Cells in culture flask were detached using 0.25 mg/ml trypsin (Gibco). PA gels were air-dried for 10 min, then droplets containing 10,000 cells were seeded in the form of monolayers on the gel surface. Monolayers were left to grow on soft of stiff gels for 5 days at 37 °C in 5 % CO_2_ to ensure their long-term priming by defined ECM stiffness. After this priming period, collagen gels were prepared. Collagen type I solution (rat tail, 4.0 mg/ml, Advanced Biomatrix) was diluted in cold media to yield a working collagen concentration of 2.3 mg/ml. This solution was adjusted to 7.5-7.8 pH using 1 M NaOH. Once this pH was reached, a sonicated solution containing 1 μm fluorescent beads (Thermo Fisher) was added to achieve a final concentration of 4μl fluorescent beads per ml of solution. This solution, containing collagen and beads, was carefully mixed to prevent bubbles, then was pipetted into the wells of a chilled glass bottom 24-well plate (Fisher Scientific). PA gels containing primed cells were carefully placed on top of this collagen solution, PA gel side down. The plate was then incubated at 37°C and 5 % CO_2_ for 45 min to facilitate crosslinking. After crosslinking this lower collagen gel layer, another layer of collagen solution with fluorescent beads was added on top of the PA gel. To facilitate crosslinking of this upper collagen layer, gels were incubated for another 45 min. Finally, media was added, with care taken to not disrupt the collagen gels. Completed gel constructs were incubated at 37°C and 5 % CO_2_ for 3 days to test the duration of the mechanical memory phenotype we might observe. Note that in this setup, cells are not trypsinized after priming, before implanting into collagen. It was necessary to devise a system in which cells are primed and then allowed to invade without detachment, because trypsinization is a harsh treatment that resets cellular state and obviates memory, as shown previously (*20*). Moreover, chemical detachment of cells while crossing the interface between two environments is unrealistic.

### Immunofluorescent imaging of cells

After 3 days of invasion, constructs were prepared for immunofluorescence imaging. Cells were fixed with 4 % paraformaldehyde for 15 min, washed with DPBS, then permeabilized with 0.3 % solution of Triton X-100 (Santa Cruz) for 10 min. Non-specific binding was blocked by 2 % bovine serum albumin (BSA) in DPBS overnight at 4°C, followed by 2 washes with DPBS. To visualize nuclei, cells were incubated with Hoechst 33258 (1:50; Thermo Fisher) for 30 min at room temperature, then washed with DPBS. For actin visualization, gels were incubated with phalloidin (1:125; Invitrogen) for 35 min and washed with DPBS. Stained gels were stored at 4°C until imaging. Fluorescent images were recorded using a laser-scanning confocal microscope (Zeiss LSSM 730; Carl Zeiss Micro Imaging, Germany) at 10×, 20×, and 40× objectives, and z-stacks were acquired at 5 μm intervals. Experiments were performed in triplicates and quadruples per well. Meanwhile, laser intensity and exposure time were kept constant to enable quantitative analysis across samples. To prevent potential biasing, the images used for analysis were randomly selected from 6-7 fields of view for each condition.

### Time-lapse microscopy

Live-cell imaging to visualize collective cell migration and fluorescent bead movement was done using a Zeiss Cell Observer microscope (10x objective, Carl Zeiss Microscopy) equipped with an incubation chamber. In each experiment, phase-contrast and tracking of 1 μm orange (540/560) fluorescent beads images were acquired to facilitate analysis of cell invasion and bead displacement, respectively. Images were collected every 1 h for 48-72 h and cells were maintained at 37 °C and 5% CO_2_ for the duration of time-lapse imaging.

### Modulation of collagen density and collagen crosslinking

To determine how primed cells interact with different collagen densities and fiber structure, collagen gels of three different modalities: 2.3mg/ml, 2.3mg/ml+crosslinked, and 3.1 mg/ml were constructed, using the same procedure described above. Collagen crosslinking was achieved by addition of Riboflavin 5’ phosphate sodium slat hydrate (Riboflavin; Sigma Aldrich) to the media-collagen solution (final concentration 0.5 mM). This collagen solution was then prepared as previously described to create the two fully crosslinked layers. After media was added, the constructs were exposed to 365 nm UV for 15 s to activate riboflavin and crosslink collagen fibers to create bundle and potentially thicker fibers in the gel.

### Spatial and temporal analyses of cell invasion

Imaris software (Bitplane) was used to create a 3D reconstruction of the invading cells in collagen utilizing confocal images and stacks. A custom batch process was developed to reconstruct cell nuclei from the Hoechst signal, record cell coordinates in the 3D space, and count number of invaded cells at given distance. A custom MATLAB code was written to calculate the distance vector of the cells in relation to the edge of the gel. To determine the rate of invasion of the primed cells into the collagen, rectangular regions of interest (RoIs) of dimensions 387 μm × 213 μm were selected within collagen and cell migration was analyzed from time-lapse videos. A custom ImageJ (https://imagej.net/Fiji) macro was developed to identify invading cells using a gaussian blur. From this macro, the area occupied by the invading cells was extracted and Microsoft Excel was used to calculate the difference between area occupied by invaded cells at time *t* and area at time *t*=0, and this difference by normalized by the width of the rectangular RoI to yield invasion distance (μm) per unit hour. To determine the advantage of stiff-primed cells over soft-primed cells, we also calculated the difference in their invasion rate, termed here as the “priming advantage”.

### Collagen deformation and accumulation analyses

The RoI approach described above was also used to track the collagen deformation rate over time. To track the fluorescent beads, particle image velocimetry (PIV) analysis was done to calculate spatiotemporal profiles of velocity magnitudes through PIVlab package in MATLAB (*34*). For each condition, PIV was performed by using three passes of 64-, 32- and 16-pixel windows to obtain the velocity field (v_i_). These values were exported as MATLAB workspace. A custom MATLAB code was used to calculate the average velocity of the beads over time to determine the amount of deformation of the collagen gel due to primed cells invasion. Collagen accumulation can be quantified by the changes to the beads fluorescent intensity since the beads are attached to collagen fibers. The bead fluorescent intensity in collagen gels was used to create kymographs by calculating the bead fluorescent intensity in five regions along the length of the rectangular ROI. The regions were defined by how close they were to the edge of the PA gel. Using ImageJ, the intensities in each region were extracted then averaged among samples and regions. Normalization of bead intensity across different regions was done against intensity at *t*=*0*. Using these values, a kymograph was plotted to display temporal and spatial changes in bead fluorescent intensity.

### Collagen relaxation analysis

To assess net tension in bulk collagen stored by the primed cells after they invaded for 3 days, trypsinization was used to cause cellular detachment from collagen to quantify differences in collagen displacements. For collagen displacement, 1 μm fluorescent beads were embedded in 2.3 mg/ml collagen. MCF10As were then allowed to invade for three days, then trypsin was added causing cells to detach from the collagen. The tracking of the bead displacement was done using a confocal microscopy. Fluorescent beads were imaged at the same laser power and exposure. These images were processed through PIV to get vector components of beads movement, which were processed in MATLAB to calculate bead displacement in the X and Z planes and plot heatmaps of bead displacement, superimposed with locations of cells (as black circles) present in collagen before trypsinization.

### Collagen visualization and OrientationJ collagen alignment analysis

To visualize collagen structure and fiber orientation, reflectance was used through a small wide wavelength that passes through the sample and then captures the back scattering of light passing through the collagen matrix allowing us to visualize collagen fiber structures. Reflection images of collagen were taken using a 20X objective, 5 μm z-stacks, and with consistent laser power. To analyze the orientation of the collagen, we used the Z-projection of images and utilized OrientationJ Fiji Plugin in combination with a batch ImageJ macro to calculated fiber orientation angles. This is paired with an R script that normalizes the angle distribution between −90°and 90° (*35, 36*). Similar analysis was done for the images with F-actin fibers.

### Correlation length analysis for bead deformation rate

To understand whether collagen bead displacement at one place in collagen is correlated with displacement elsewhere, we calculated correlation length, which has conventionally been used for collective cell migration. Using a custom MATLAB code that we previously developed for cells (*37*), correlation coefficients were averaged over all directions and plotted as function of the distance between the index position, the distance between the two positions, and the width of the annulus. The correlation length is defined as the slope of the negative exponential fit. This was calculated in MATLAB, then averaged per time point and fitted in Prism 8. These correlation lengths were then averaged across minimum 3 biological replicates to get a mean correlation length value and plotted as histograms in Prism 8 for both conditions.

### Fiber coherency

To determine level of coherency of collagen or F-actin fibers, OrientationJ plugin in ImageJ was used. Randomly 9-10 RoIs were selected and coherency value was calculated using collagen reflectance or F-actin images, applicable. Coherency values were averaged across samples for each condition.

### Collagen bundling analysis

To quantify collagen bundling after 3 days of priming, we extracted ROIs of similar sizes and developed a custom macro in ImageJ to determine the degree of fiber bundling seen in reflectance images. To analyze collagen bundling, first the images were processed using ImageJ in these steps: mean intensity within a 960μm × 960μm ROI is measured; this mean was subtracted from the ROI; images are converted to binary; final raw intensity yields the degree of bundling of collagen fibers. To normalize bundling across images, raw intensity values were divided by 255 to get the number of pixels, which were divided by the area of the ROI.

### Measuring pore size in collagen

Utilizing reflectance images of each collagen density, an ImageJ macro was developed to extract the Feret diameter from a stack of images. The macro selects the stack placing a Gaussian blur on the image. Then, the threshold is adjusted to highlight the empty spaces found between fibers; these spaces represent pores within the collagen matrix. This threshold value is applied to create a binary image and Feret diameter, representing pore size is measured for all Z-slices.

### Pharmacological inhibition studies

To understand the effects of contractility and collagen crosslinking, inhibitors were used during live imaging to quantify their effect. For inhibition of myosin II, 20mM Blebbistatin (Sigma Aldrich) was added after 30 hours of invasion. For LOX inhibition, 10mM β-Aminopropionitrile (BAPN; Sigma Aldrich), was added 10 minutes before live imaging.

### qPCR of primed cells

MCF10a and shYAP MCF10a were left to prime for 5 days on stiff and soft PA gels. The RNA was isolated from the cells using the Qiagen’s RNeasy kit following manufacturer instructions. The RNA quality and quantity were measured using Nano dropper (Thermo Fisher) then standardized to 200 ug/ml and converted to cDNA using the C1000 Thermocycler (Bio-Rad). A volume of cDNA was added to each well of the 96 well plate that contained 10 μl of a solution of TaqMan Fast Advanced Master Mix (Applied Biosystems), nuclease free water (Invitrogen), and TaqMan primers (Thermo Fisher). We used B2M (Assay ID: Hs00187842) as our housekeeping gene and ran reactions for Rac (Assay ID: Hs01902432), RhoA (Assay ID: Hs00357608), and Lox (Assay ID: Hs00942480) using QuantStudio real time PCR in triplicates for an N=6. Expressions of each gene were normalized to the housekeeping gene. The relative expressions were calculated from the comparison of the difference between in Ct between target genes and the reference gene. This fold change was done in comparison of stiff to soft. For list and sequences of used oligonucleotides refer to Table 1.

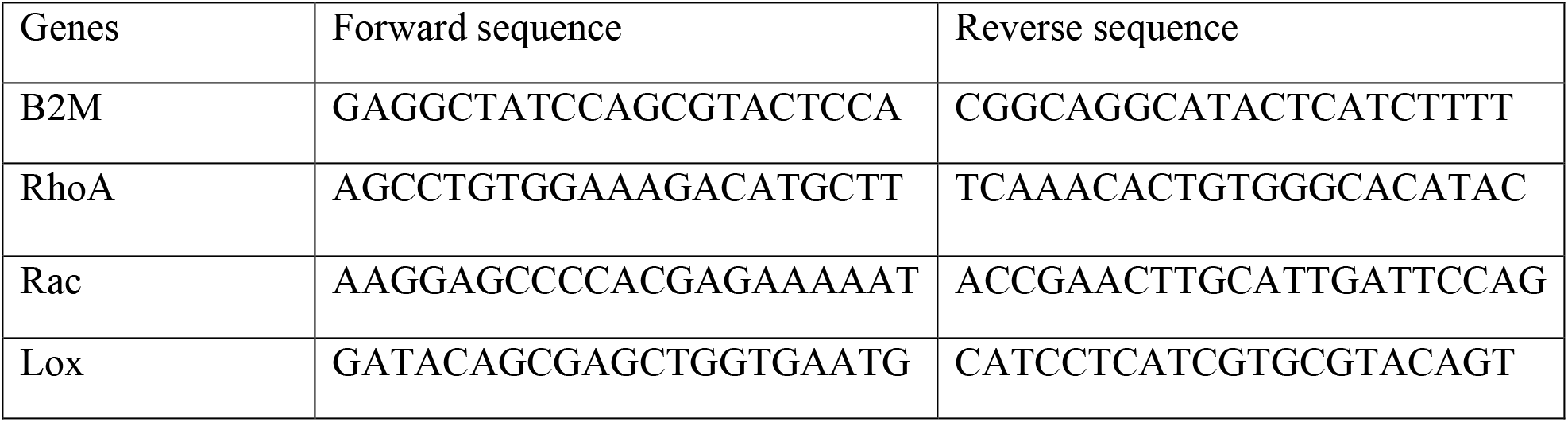

### Laser ablation of collagen

To ablate collagen fibers, we used an Andor Micropoint Laser Ablation system connected to the Zeiss Cell Observer system and controlled with Andor i8 software (Andor). A straight line was ablated in the collagen before invasion began, creating a cut approximately 150 μm away, parallel from the edge of the gel. The cut was made on the Z-plane where the cells appeared in focus. Bead tracking for laser ablation was done using PIV. Fixed imaging of the cut collagen and invaded cells was performed as described above.

### Mechanical characterization of collagen stiffness for different concentrations

To quantify Young’s Modulus of collagen of different compositions, atomic force microscopy (AFM) was performed. Collagen stiffness was measured using a Bruker BioScope Resolve Bio-atomic force microscope (AFM; Bruker). AFM probes were custom-made by Nanoscope to have a 4.5 μm bead attached to the cantilever with a nominal stiffness 0.01 N/m. Elastic moduli were analyzed from collected force curves using the Hertz model (*38*).

### Scanning electron microscopy

Scanning electron microcopy was performed to visualize the collagen alignment and cells as they invade into the collagen. The *in vitro* PA-collagen gel system with invaded cells, after 5 days of priming and 3 days of invasion described above, was fixed using 2.5 % glutaraldehyde that has been mixed with a solution of 2% paraformaldehyde in 0.15 M cacodylate buffer at a pH of 7.4 with 2 mM calcium chloride. Once warmed, cell media was removed, and a fixative was added and incubated at 35 °C and 5 % CO_2_ for 15 min. Samples were removed from the incubator and placed in a shaker overnight at room temperature. Images were acquired using a high-resolution scanning electron microscope (Zeiss Merlin FE-SEM) at Washington University Center for Cellular Imaging (WUCCI).

### Statistics and reproducibility

All data represent at least 4 replicates from separate experiments. All bar graphs are presented as mean ± standard error. All box and whiskey plots are mid-line represent mean and 90%-10% whiskeys. Similar sample size and statistical tests were used for each experiment, and these are indicated on figure legends. Statistical significance was calculated majority with test for pairwise comparisons and one-way analysis of variance (ANOVA), unless specified otherwise. All statistical analyses were performed in Prism.

## Computational Model

### Overview: modeling memory-dependent invasion

Our experimental findings connect cellular memory to direct mechanosensing along with an active ECM remodeling feedback. Thus, several intercoupled cellular and extracellular processes, some mechanical and some biochemical, give rise to memory-dependent invasion. Here, we develop a theoretical model and a computational framework to better understand these complex processes, which cannot be explained by existing models of cell migration resulting from sensing current mechanical environments.

### Cell migration as an energy minimization problem

We recall that cell migration has traditionally been understood as three basic steps (*39*): (a) extension of frontward protrusions attempting to propel the cell forward; (b) formation of cell-ECM focal adhesions that provide traction and resistance against motion; (c) generation of cytoskeletal contractile forces that break adhesions to reduce resistance and enable net cell translocation. How these three processes generate cell migration can be interpreted as an energy minimization problem. From a resting state, biochemical signaling for actin polymerization gives rise to protrusions, and this protrusive energy is minimized by anchoring protrusions via adhesion formation. Mechanotransductive signaling gives rise to actin-myosin contractile forces that pull of focal adhesion. As a result, there is potential energy stored in stretched focal adhesions, which can be minimized by breaking adhesions. As such, the forward translocation of the cell can be understood as minimizing energy fluctuations due to dynamic evolution of protrusions, contraction, and adhesions. Cell migration would not occur if energy does not rise due to protrusions or contraction-based stretching of adhesions. We formulate a net energy balance (Fig. S12A) as:

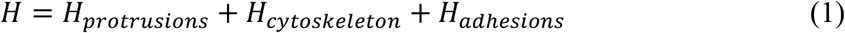

Here, *H_protrusions_* is the energy associated stable protrusions, *H_cytoskeleton_* is the energy associated with cell body properties like supracellular contractility and cell rigidity that helps maintain cell shape, and *H_adhesions_* is the energy associated with cell contacts (combined cell-cell and cell-ECM adhesions in this case). In 3D environments, cells experience resistance not only from adhesions but also drag due to bulk crowding of ECM proteins. For net cell movement to occur, the change in net energy Δ*H* should be less than zero. Unlike 2D substrates, cells can align and remodel 3D fibrous matrices (*40*), which is an additional energy term that is not accounted for in the energy consideration described above (Eq. 1). We propose that collagen remodeling acts to lower the energy barrier for invasion (Fig. S12B). We implement this process through Δ*H_protrusions_*, because net stable protrusions must overcome ECM barriers in 3D, written as:

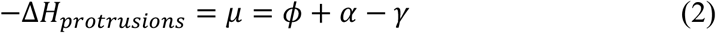

This formulation is analogous to previous models of 3D cell invasion that calculate net cell movement as a sum of forces due to contractility, protrusion, and drag (*41*). Here, our key addition is the active feedback between contractile forces and ECM resistance via force-based ECM remodeling. In Eq. 2, the direct mechanosensing signal from the ECM (*ϕ*), collagen remodeling (*α*), and resistance via ECM (*γ*) together result in net protrusive potential *μ,* which lowers Δ*H_protrusions_* in the net energy balance (Eq. 1). This conceptual framework of energy minimization causing cell invasion in 3D environments is somewhat independent of computational methods, in principle, and can be implemented in agent-based, element-based, lattice-based, or vertex modeling methodologies. Here, we utilize the Cellular Potts method (*42, 43*) to implement this model due to its several advantages – computational efficiency, description of the ECM as a dynamic energy field, and relatively straightforward cell-cell and cell-ECM interactions.

### Defining adhesion and bulk cell energies

Cellular Potts model (*43*) represents a space as a discrete collection of lattice points – pixels on a 2D grid or voxels in a 3D grid. We model a collection of biological cells by attaching to each lattice point (*i; j*) of a square lattice a label σ_*ij*_, which identifies the corresponding cell, and a label *τ*(σ_*ij*_), which identifies cell type. Adjacent lattice sites are defined to lie within the same cell if they have the same value of σ_*ij*_. System evolves by the random movement of individual pixels that move according to transition probabilities based on Monte Carlo simulations based on the energy criterion described above (*44*). At each Monte Carlo Step, two neighboring pixels are chosen randomly, with one as source pixel and the other as target pixel. If both pixels belong to the same cell (*i. e*., σ(*source*) = σ(*target*)), then no changes are made to the lattice. Otherwise, the source pixel attempts to occupy the target pixel based on Monte Carlo acceptance probability, which is calculated from the difference in total system energy. The total system energy associated with the configuration, before and after the move, is defined as per Eq. 1. We provide specific definitions for each term in Eq. 1, as following:

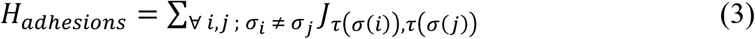

Here, *J*_τ_(*_σ_*(*_i_*)),*_τ_*(*_σ_*(*_j_*)) represents contact energy for the two cell types in contact and (*τ*(σ(*i*))*τ*(σ(*j*)) represents contribution from the total energy due to cell-cell adhesions.

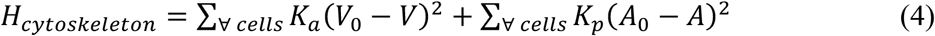

In *H_cytoskeleton_*, two terms represent contributions from bulk elasticity of the cell and cell-surface contractility, respectively. *K_a_* and *K_p_* are constants for bulk elasticity and contractility, respectively. *V*_0_ and *A*_0_ are target volume and surface area that the cell has in isolation. After calculating energy of system before (*H_i_*) and after (*H_f_*) the copy attempt will always be successful if *H_f_* < *H_i_*, *i. e*. Δ*H* < 0. If Δ*H* ≥ 0, the copy attempt is accepted with a probability of 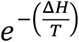, *T* = 20. Higher values of *T* would tend to accept more unfavorable copy attempt.

### Defining protrusive potential to model directed migration into collagen

Earlier, we described *H_protrusions_* as the net protrusive energy for cell migration. For net motion at any given time step, Δ*H_protrusions_* must be less than zero. In other words, –Δ*H_protrusions_* = *μ* > 0 will propel migration into the collagen; if *μ* ≤ 0, there is no bias due to protrusions to migrate in a preferred direction. As cells enter a 3D matrix of normalized density 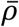, resistance (*γ*) due to matrix density must be overcome to enable invasion, as the cells sense this new environment and adapt to it (*ϕ*). We propose that cells overcome this resistance via collagen remodeling (*α*). Thus, net protrusions *μ* is defined as:

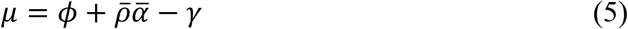

Here, *ϕ* is the direct mechanosensing signal from the current ECM. We note that *ϕ* is an abstract quantity that isolates signal due to direct mechanosensing alone, which cannot be measured, because the measurable cell state (*μ*) is the net sum of its real context – a combination of memory, ECM resistance, and remodeling. With increasing collagen density, the pore size in collagen matrices reduces, which increases resistance. Simultaneously, the rise in ligand density has been shown to give rise to biphasic cell migration (*45, 46*). Thus, we model the ECM resistance *γ* proportional to square of density (*ρ*): *γ* = *kρ*^2^, where *k* = 3 × 10^−2^ *ml*^2^*mg*^−2^ is the proportionality constant. The term 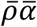 in Eq. 5 represents collagen remodeling (*21*), which lowers energy barrier posed by resistance *γ*. Here, *ϕ* and *α* evolve over time according to changes in the ECM and cell state.

### Encoding matrix memory – mechanosensing and mechanical memory combine to remodel collagen

As mechanically primed cells enter collagen, they apply forces on collagen fibers. Cellular forces are understood to arise from the net mechanoactivation state of the cell. According to previous models of cell migration, rapid mechanotransduction due to focal adhesion signaling enables direct mechanosensing of current ECM conditions (stiffness, architecture). Based on our experimental findings and previous results of cell state regulation from mechanical memory (*47, 48*), we update this model by formulating collagen remodeling *α* resulting from a combination of memory *ψ* and direct mechanosensing *ϕ* (Fig. S12C):

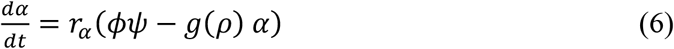

where *r_α_* = 10^−2^ *MCS*^−1^ = 1.7 × 10^−4^ *s*^−1^. The first term represents contribution to remodeling from two the cellular signals – mechanical memory *ψ* of the past ECM and direct mechanosensing *φ* of the current ECM (Fig. 3A, Fig. S12C). Mechanically, it should be easier to pull and remodel softer ECM compared to rigid ECM, thus *g*(*ρ*) represents the bulk stiffness of the collagen matrix and is defined as *g*(*ρ*) = (*ρ*/3.7)^2^. We sum *α* over the entire simulation space to obtain net remodeling 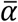 (normalized by the simulation space area), which represents the average tension buildup in the ECM as cells remodel it.

### Differential kinetics of cellular memory and direct mechanosensing

As noted above, the mechanosensing signal *φ* represents direct sensing of the current ECM and rapid mechanotransductive response that would predict quick adaptation, written as (Fig. S12C):

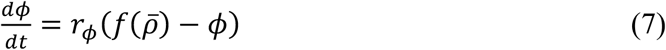

Here 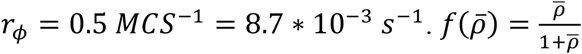 is the mechanosensitive signal due to collagen stiffness and pore size, both of which affect cell migration in 3D (*49, 50*) 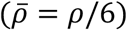. As collagen density increases, the mechanosensing signal increases, and eventually saturates (*51–53*).

In contrast to direct mechanosensing, mechanical memory *ψ* is slow to decay and adapt to the new environment, consistent with previous experimental findings of YAP-based memory regulation (*9, 20*) such that nuclear YAP localization continues to retain cellular mechanoactivation, written as:

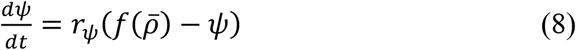

Here, the governing rate *r_ψ_* for memory kinetics in modeled as a switch-like system such that the rate is low (~0) in the beginning when cells start invading into collagen and approaches *r_ψ_* (same as that for *φ* in Eq. 7) in about 2 days:

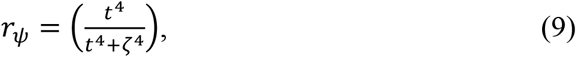

where *ζ* = 800 *MCS*~12 *H*. Lower value of *ζ* reduce memory by enhancing adaptation of *ψ* to the current collagen matrix.

### Simulation details

We utilize CompuCell3D to solve the system of differential equations described above over a defined field (150 pixels × 75 pixels, 1px = 2*μm*). At each time step, *ϕ, ψ, μ, γ* are solved for each cell, whereas *α* is also solved over space. Total simulation time is 3000 MCS, which are equivalent to 48 hours. At *t* = 0 *MCS, ϕ* = *ψ* = 1 for stiff-primed cells, and *ϕ* = *ψ* = 0.3 for soft-primed cells, with *α* = 0 everywhere. Cell monolayer is defined as a flat collection of cells of diameter= 10*μm*. Here, we assume that the collagen matrix is in a 3D space and show simulation for a given 2D plane of cell invasion.

## Acknowledgements

The authors acknowledge all members of the A.P. laboratory for discussions and feedback on this work; C. Walter for technical assistance with AFM; G.D. Longmore for providing cells; and Washington University Center for Cellular Imaging (WUCCI) for scanning electron microscopy. We acknowledge financial support from following sources: National Institutes of Health grant R35GM128764 (to A.P.) and National Science Foundation, Science and Technology Centers, Center for Engineering MechanoBiology grant CMMI:154857 (to A.P.).

## Author Contributions

Conceptualization: AP

Methodology: JA, BS, AP

Investigation: JA, JM, AP

Visualization: JA, JM, AP

Formal Analysis: JA, JM, YLL

Funding acquisition: AP

Project administration: AP

Supervision: AP

Writing – original draft: JA, JM, AP

Writing – review & editing: AP

## Data and materials availability

All data are available in the main text or the supplementary materials

## References

1. A. Shellard, A. Szabó, X. Trepat, R. Mayor, Supracellular contraction at the rear of neural crest cell groups drives collective chemotaxis. Science 362, 339–343 (2018).

2. M. J. Paszek, N. Zahir, K. R. Johnson, J. N. Lakins, G. I. Rozenberg, A. Gefen, C. A. Reinhart-King, S. S. Margulies, M. Dembo, D. Boettiger, D. A. Hammer, V. M. Weaver, Tensional homeostasis and the malignant phenotype. Cancer Cell 8, 241–254 (2005).

3. I. Acerbi, L. Cassereau, I. Dean, Q. Shi, A. Au, C. Park, Y. Y. Chen, J. Liphardt, E. S. Hwang, V. M. Weaver, Human breast cancer invasion and aggression correlates with ECM stiffening and immune cell infiltration. Integr Biol (Camb) 7, 1120–1134 (2015).

4. R. J. Pelham, Y.-l. Wang, Cell locomotion and focal adhesions are regulated by substrate flexibility. Proceedings of the National Academy of Sciences, USA 94, 13661–13665 (1997).

5. C. X. Li, N. P. Talele, S. Boo, A. Koehler, E. Knee-Walden, J. L. Balestrini, P. Speight, A. Kapus, B. Hinz, MicroRNA-21 preserves the fibrotic mechanical memory of mesenchymal stem cells. Nat Mater 16, 379–389 (2017).

6. M. R. Ng, A. Besser, G. Danuser, J. S. Brugge, Substrate stiffness regulates cadherin-dependent collective migration through myosin-II contractility. J Cell Biol 199, 545–563 (2012).

7. R. Sunyer, V. Conte, J. Escribano, A. Elosegui-Artola, A. Labernadie, L. Valon, D. Navajas, J. M. García-Aznar, J. J. Muñoz, P. Roca-Cusachs, X. Trepat, Collective cell durotaxis emerges from long-range intercellular force transmission. Science 353, 1157–1161 (2016).

8. C. C. Price, J. Mathur, J. D. Boerckel, A. Pathak, V. B. Shenoy, Dynamic Self-Reinforcement of Gene Expression Determines Acquisition of Cellular Mechanical Memory. Biophysical Journal, (2021).

9. S. Nasrollahi, C. Walter, A. J. Loza, G. V. Schimizzi, G. D. Longmore, A. Pathak, Past matrix stiffness primes epithelial cells and regulates their future collective migration through a mechanical memory. Biomaterials 146, 146–155 (2017).

10. M. A. Wozniak, R. Desai, P. A. Solski, C. J. Der, P. J. Keely, ROCK-generated contractility regulates breast epithelial cell differentiation in response to the physical properties of a three-dimensional collagen matrix. Journal of Cell Biology 163, 583–595 (2003).

11. E. Cukierman, R. Pankov, D. R. Stevens, K. M. Yamada, Taking Cell-Matrix Adhesions to the Third Dimension. Science 294, 1708–1712 (2001).

12. O. Ilina, P. G. Gritsenko, S. Syga, J. x. F. C. r. Lippoldt, C. A. M. L. Porta, O. Chepizhko, S. Grosser, M. Vullings, G.-J. Bakker, J. x. F. r. S. x000Df, P. Bult, S. Zapperi, J. A. K. x. E. s, A. Deutsch, P. Friedl, Cell–cell adhesion and 3D matrix confinement determine jamming transitions in breast cancer invasion. Nature Cell Biology 17, 1–33 (2020).

13. P. P. Provenzano, D. R. Inman, K. W. Eliceiri, S. M. Trier, P. J. Keely, Contact guidance mediated three-dimensional cell migration is regulated by Rho/ROCK-dependent matrix reorganization. Biophys J 95, 5374–5384 (2008).

14. C. M. Kraning-Rush, S. P. Carey, M. C. Lampi, C. A. Reinhart-King, Microfabricated collagen tracks facilitate single cell metastatic invasion in 3D. Integrative Biology 5, 606 (2013).

15. B. R. Seo, X. Chen, L. Ling, Y. H. Song, A. A. Shimpi, S. Choi, J. Gonzalez, J. Sapudom, K. Wang, R. C. A. Eguiluz, D. Gourdon, V. B. Shenoy, C. Fischbach, Collagen microarchitecture mechanically controls myofibroblast differentiation. Proceedings of the National Academy of Sciences 117, 1–12 (2020).

16. M. Chrzanowska-Wodnicka, K. Burridge, Rho-stimulated contractility drives the formation of stress fibers and focal adhesions. J Cell Biol 133, 1403–1415 (1996).

17. M. H. Zaman, R. D. Kamm, P. Matsudaira, D. A. Lauffenburger, Computational model for cell migration in three-dimensional matrices. Biophys J 89, 1389–1397 (2005).

18. D. A. Lauffenburger, A. F. Horwitz, Cell Migration: A Physically Integrated Molecular Process. Cell 84, 359–369 (1996).

19. H. Ahmadzadeh, M. R. Webster, R. Behera, A. M. J. Valencia, D. Wirtz, A. T. Weeraratna, V. B. Shenoy, Modeling the two-way feedback between contractility and matrix realignment reveals a nonlinear mode of cancer cell invasion. Proceedings of the National Academy of Sciences 114, E1617–E1626 (2017).

20. C. Yang, M. W. Tibbitt, L. Basta, K. S. Anseth, Mechanical memory and dosing influence stem cell fate. Nat Mater 13, 645–652 (2014).

21. K. Wolf, Y. I. Wu, Y. Liu, J. Geiger, E. Tam, C. Overall, M. S. Stack, P. Friedl, Multi-step pericellular proteolysis controls the transition from individual to collective cancer cell invasion. Nat Cell Biol 9, 893–904 (2007).

22. P. Grunert, B. H. Borde, S. B. Towne, Y. Moriguchi, K. D. Hudson, L. J. Bonassar, R. Hartl, Riboflavin crosslinked high-density collagen gel for the repair of annular defects in intervertebral discs: An in vivo study. Acta Biomater 26, 215–224 (2015).

23. A. G. Gouldin, M. E. Brown, J. L. Puetzer, An inducible model for unraveling the effects of advanced glycation end-product accumulation in aging connective tissues. Connect Tissue Res, 1–19 (2021).

24. C.-M. Lo, H.-B. Wang, M. Dembo, Y.-l. Wang, Cell Movement Is Guided by the Rigidity of the Substrate. Biophysical Journal 79, 144–152 (2000).

25. P. Friedl, K. Wolf, Plasticity of cell migration: a multiscale tuning model. J. Cell Biol. 188, 11–19 (2010).

26. C. Beadle, M. C. Assanah, P. Monzo, R. Vallee, S. S. Rosenfeld, P. Canoll, The Role of Myosin II in Glioma Invasion of the Brain. Mol Biol Cell 19, 3357–3368 (2008).

27. A. Shellard, R. Mayor, Collective durotaxis along a self-generated stiffness gradient in vivo. Nature, 1–5 (2021).

28. S. B. Klein, What memory is. Wires Cogn Sci 6, 1–38 (2015).

29. K. A. U. Gonzales, L. Polak, I. Matos, M. T. Tierney, A. Gola, E. Wong, N. R. Infarinato, M. Nikolova, S. Luo, S. Liu, J. S. S. Novak, K. Lay, H. A. Pasolli, E. Fuchs, Stem cells expand potency and alter tissue fitness by accumulating diverse epigenetic memories. Science 374, eabh2444 (2021).

30. B. Seelbinder, S. Ghosh, S. E. Schneider, A. K. Scott, A. G. Berman, C. J. Goergen, K. B. Margulies, K. C. Bedi, Jr., E. Casas, A. R. Swearingen, J. Brumbaugh, S. Calve, C. P. Neu, Nuclear deformation guides chromatin reorganization in cardiac development and disease. Nat Biomed Eng 5, 1500–1516 (2021).

31. A. R. Killaars, J. C. Grim, C. J. Walker, E. A. Hushka, T. E. Brown, K. S. Anseth, Extended Exposure to Stiff Microenvironments Leads to Persistent Chromatin Remodeling in Human Mesenchymal Stem Cells. Advanced Science 17, 1801483–1801413 (2018).

32. R. S. Stowers, A. Shcherbina, J. Israeli, J. J. Gruber, J. Chang, S. Nam, A. Rabiee, M. N. Teruel, M. P. Snyder, A. Kundaje, O. Chaudhuri, Matrix stiffness induces a tumorigenic phenotype in mammary epithelium through changes in chromatin accessibility. Nature Biomedical Engineering 139, 1–14 (2019).

33. A. J. Loza, S. Koride, G. V. Schimizzi, B. Li, S. X. Sun, G. D. Longmore, Cell density and actomyosin contractility control the organization of migrating collectives within an epithelium. Mol Biol Cell 27, 3459–3470 (2016).

34. W. Thielicke, E. J. Stamhuis, PIVlab – Towards User-friendly, Affordable and Accurate Digital Particle Image Velocimetry in MATLAB. Journal of Open Research Software 2, 1202 (2014).

35. A. Kaur, B. L. Ecker, S. M. Douglass, C. H. Kugel, M. R. Webster, F. V. Almeida, R. Somasundaram, J. Hayden, E. Ban, H. Ahmadzadeh, J. Franco-Barraza, N. Shah, I. A. Mellis, F. Keeney, A. Kossenkov, H.-Y. Tang, X. Yin, Q. Liu, X. Xu, M. Fane, P. Brafford, M. Herlyn, D. W. Speicher, J. A. Wargo, M. T. Tetzlaff, L. E. Haydu, A. Raj, V. Shenoy, E. Cukierman, A. T. Weeraratna, Remodeling of the Collagen Matrix in Aging Skin Promotes Melanoma Metastasis and Affects Immune Cell Motility. Cancer Discovery 9, 64–81 (2019).

36. J. Franco-Barraza, D. A. Beacham, M. D. Amatangelo, E. Cukierman, Preparation of Extracellular Matrices Produced by Cultured and Primary Fibroblasts. Curr Protoc Cell Biol 71, 10 19 11–10 19 34 (2016).

37. B. Sarker, A. Bagchi, C. Walter, J. Almeida, A. Pathak, Longer collagen fibers trigger multicellular streaming on soft substrates via enhanced forces and cell–cell cooperation. Journal of Cell Science 132, jcs226753 (2019).

38. J. L. MacKay, S. Kumar, Measuring the elastic properties of living cells with atomic force microscopy indentation. Methods Mol Biol 931, 313–329 (2013).

39. D. A. Lauffenburger, A. F. Horwitz, Cell migration: a physically integrated molecular process. Cell 84, 359–369 (1996).

40. B. R. Seo, X. Chen, L. Ling, Y. H. Song, A. A. Shimpi, S. Choi, J. Gonzalez, J. Sapudom, K. Wang, R. C. Andresen Eguiluz, D. Gourdon, V. B. Shenoy, C. Fischbach, Collagen microarchitecture mechanically controls myofibroblast differentiation. Proc Natl Acad Sci U S A 117, 11387–11398 (2020).

41. M. H. Zaman, R. D. Kamm, P. Matsudaira, D. A. Lauffenburger, Computational model for cell migration in three-dimensional matrices. Biophys J 89, 1389–1397 (2005).

42. M. H. Swat, G. L. Thomas, J. M. Belmonte, A. Shirinifard, D. Hmeljak, J. A. Glazier, Multi-Scale Modeling of Tissues Using CompuCell3D (Methods in Cell Biology, Elsevier Inc., 2012), vol. 110.

43. F. Graner, J. A. Glazier, Simulation of biological cell sorting using a two-dimensional extended Potts model. Phys Rev Lett 69, 2013–2016 (1992).

44. M. H. Swat, G. L. Thomas, J. M. Belmonte, A. Shirinifard, D. Hmeljak, J. A. Glazier, Multi-scale modeling of tissues using CompuCell3D. Methods Cell Biol 110, 325–366 (2012).

45. M. H. Zaman, L. M. Trapani, A. L. Sieminski, D. Mackellar, H. Gong, R. D. Kamm, A. Wells, D. A. Lauffenburger, P. Matsudaira, Migration of tumor cells in 3D matrices is governed by matrix stiffness along with cell-matrix adhesion and proteolysis. Proc Natl Acad Sci U S A 103, 10889–10894 (2006).

46. P. A. DiMilla, K. Barbee, D. A. Lauffenburger, Mathematical model for the effects of adhesion and mechanics on cell migration speed. Biophys J 60, 15–37 (1991).

47. S. Nasrollahi, C. Walter, A. J. Loza, G. V. Schimizzi, G. D. Longmore, A. Pathak, Past matrix stiffness primes epithelial cells and regulates their future collective migration through a mechanical memory. Biomaterials 146, 146–155 (2017).

48. A. W. Watson, A. D. Grant, S. S. Parker, S. Hill, M. B. Whalen, J. Chakrabarti, M. W. Harman, M. R. Roman, B. L. Forte, C. C. Gowan, R. Castro-Portuguez, L. K. Stolze, C. Franck, D. A. Cusanovich, Y. Zavros, M. Padi, C. E. Romanoski, G. Mouneimne, Breast tumor stiffness instructs bone metastasis via maintenance of mechanical conditioning. Cell Rep 35, 109293 (2021).

49. P. Friedl, K. Wolf, Plasticity of cell migration: a multiscale tuning model. J Cell Biol 188, 11–19 (2010).

50. C. M. Lo, H. B. Wang, M. Dembo, Y. L. Wang, Cell movement is guided by the rigidity of the substrate. Biophys J 79, 144–152 (2000).

51. A. Saez, A. Buguin, P. Silberzan, B. Ladoux, Is the mechanical activity of epithelial cells controlled by deformations or forces? Biophys J 89, L52–54 (2005).

52. J. P. Califano, C. A. Reinhart-King, Substrate Stiffness and Cell Area Predict Cellular Traction Stresses in Single Cells and Cells in Contact. Cell Mol Bioeng 3, 68–75 (2010).

53. S. J. Han, K. S. Bielawski, L. H. Ting, M. L. Rodriguez, N. J. Sniadecki, Decoupling substrate stiffness, spread area, and micropost density: a close spatial relationship between traction forces and focal adhesions. Biophys J 103, 640–648 (2012).

