## Supplementary Materials for "Mechanically primed cells transfer memory to fibrous matrices for persistent invasion"

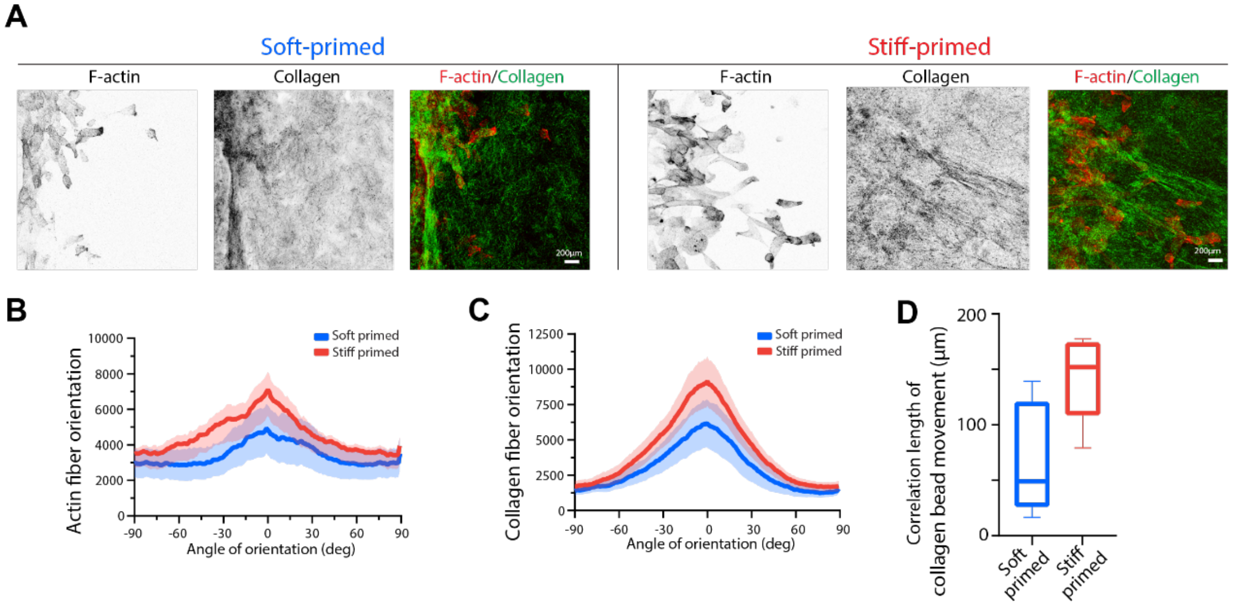


**Fig. S1. Images and analyses of collagen and actin fibers.** (**A**) Split channel images of collagen reflectance and F-actin immunofluorescence corresponding to merged images shown in Fig. 1 for primed cell invasion. Scale bar, 200µm. (**B**) Alignment distribution F-actin fibers (mean ± SEM), ranging from -90 to 90 degrees, where higher peaks in stiff primed cells (red) indicate higher number of aligned fibers compared to soft primed cells(blue). *N=8*. **(C)** Alignment distribution of collagen (mean ± SEM), ranging from -90 to 90 degrees, where higher peaks in stiff primed cells (red) indicate higher number of aligned fibers compared to soft primed cells(blue). *N=8*. **(D)** Correlation length for collagen bead displacements (from PIV analyses) caused by soft and stiff primed cells, which shows that collagen deformation due to stiff primed cells is correlated over longer distances (average distance around 150µm). *N=8*.

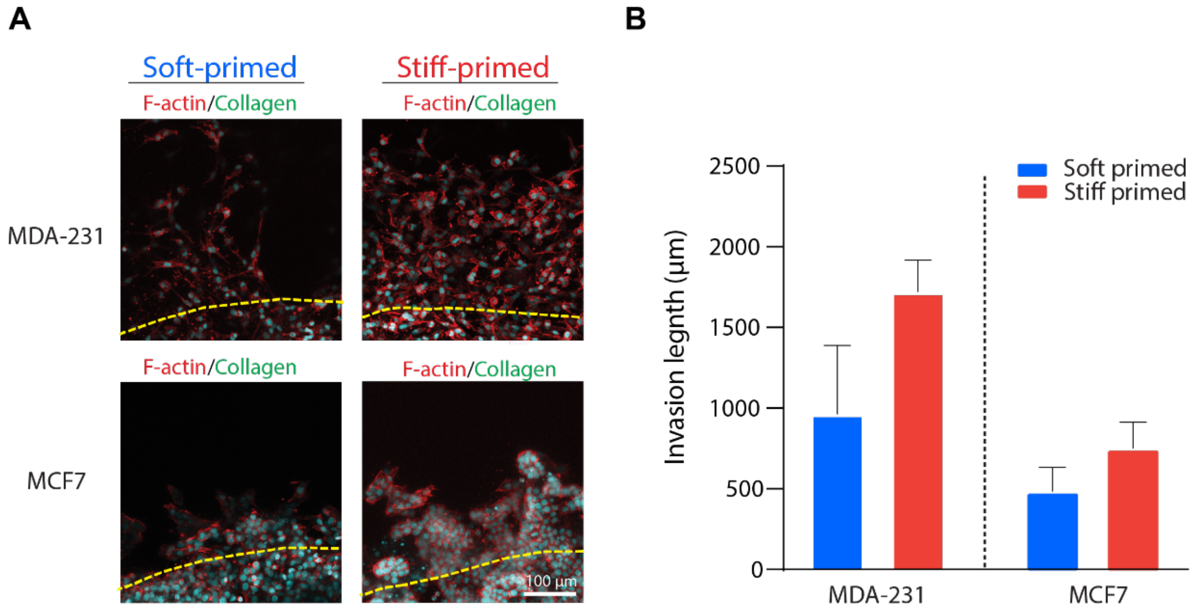


**Fig. S2. Priming-dependent invasion of MCF7 and MDA-231 cells.** (**A**) Immunofluorescence images of MCF7 and MDA-231 invaded into 2.3 mg/ml collagen after 5 days of soft or stiff priming followed by 3 days of invasion; F-actin (red) and nuclei (cyan). Yellow dotted line show the edge of the PA gel. Scale bar, 100µm. (**B**) Average length of cell invasion relative to the edge of the PA gel, which shows higher invasion by stiff-primed cells compared to soft-primed cells for both MCF7 and MDA-231 cell types. *N=6.*

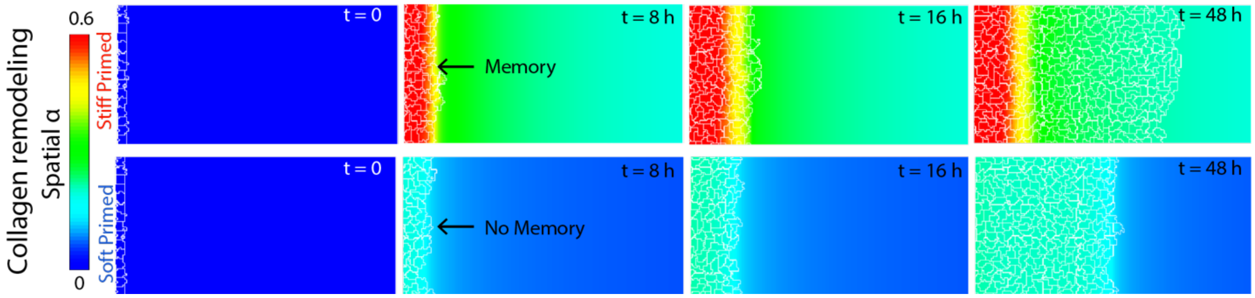


**Fig. S3. Temporal evolution of collagen remodeling for stiff primed and soft primed cells invading into collagen.** Snapshot of collagen remodeling at $t = 0,8,16\ and\ 48\ hours$ for stiff primed and soft primed cells entering collagen ($\rho = 2.3\ mg/ml$).

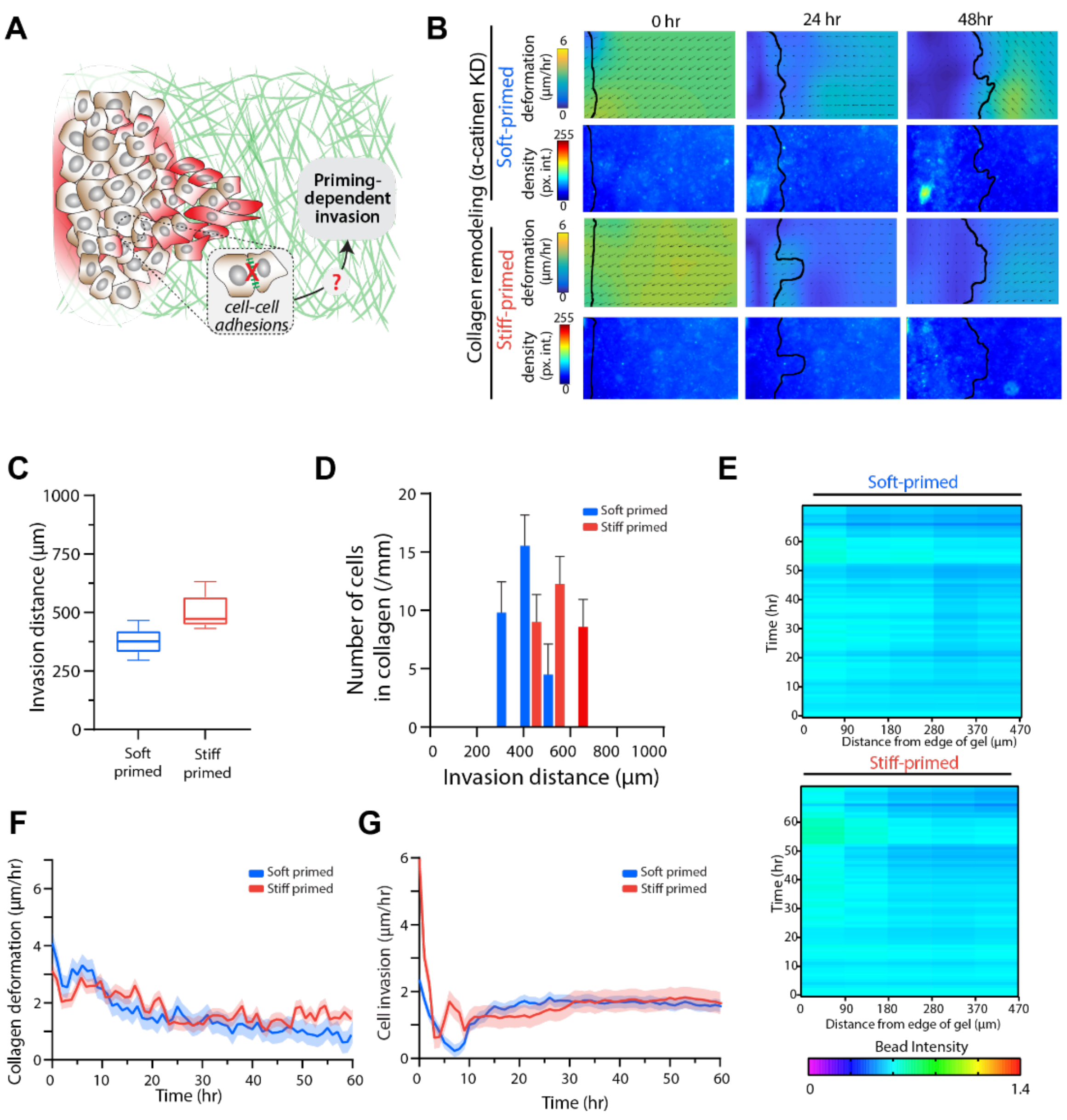


**Fig. S4. α-catenin depletion prevents coordinated collective cell invasion and disrupts priming-dependent invasion.** (**A**) Schematic posing the question of how cell-cell adhesions could disrupt cell-cell coordination and role of collective forces in matrix remodeling and invasion. (**B**) Heatmaps of collagen deformation (PIV vectors) and collagen bead intensity caused by α-catenin-KD soft or stiff primed cells (black outline annotates the cell invasion front). (**C**) Average invasion distance for soft and stiff primed cells. *N*= *5* (**D**) Histogram distribution of number of cells invaded relative to distance from the PA gel. *N*= *5*. (**E**) Kymographs of collagen bead intensity over time of invasion and distance from the PA gel for soft and stiff primed cells. *N*=*5*. (**F**) Rate of collagen deformation (mean ± SEM) and (**G**) cell invasion (mean ± SEM) over time caused by soft or stiff primed α-catenin-KD cells.

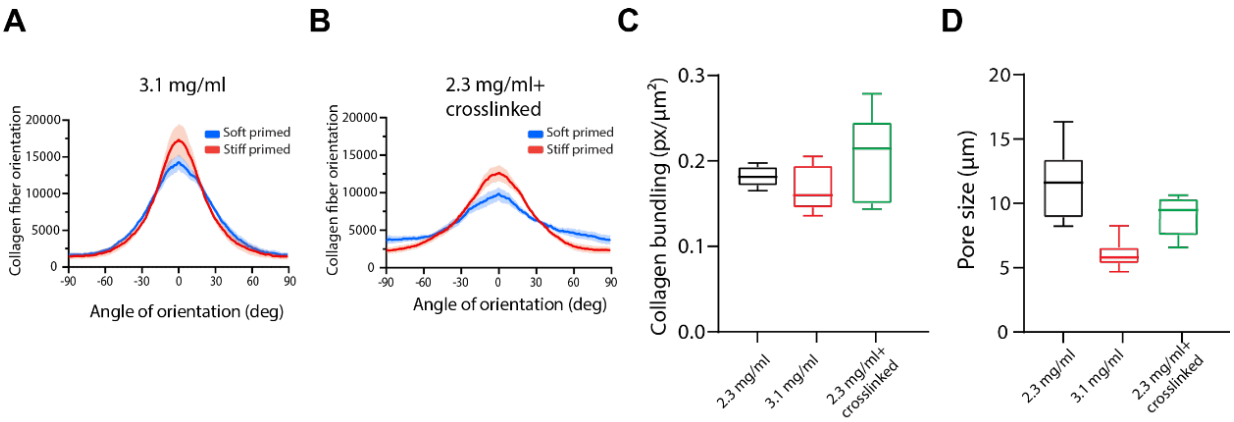


**Fig. S5. Priming-dependent matrix remodeling in dense or crosslinked collagen. (A)** Alignment distribution of collagen fibers in 3.1 mg/ml collagen, where stiff primed cells (red) and soft primed (blue) are compared showing higher collagen alignment for stiff primed cells. *N=6*. **(B)** Alignment distribution of collagen fibers in 2.3mg/ml crosslinked collagen, where stiff primed cells (red) and soft primed (blue) are compared showing higher collagen alignment for stiff primed cells. *N=6*. **(C)** Average collagen bundling after 3 days on invasion of different collagen densities caused by stiff-primed cells. *N= 8*. **(D)** Average pore size of the control (black), dense collagen (red), and crosslinked collagen (green). *N= 8*.

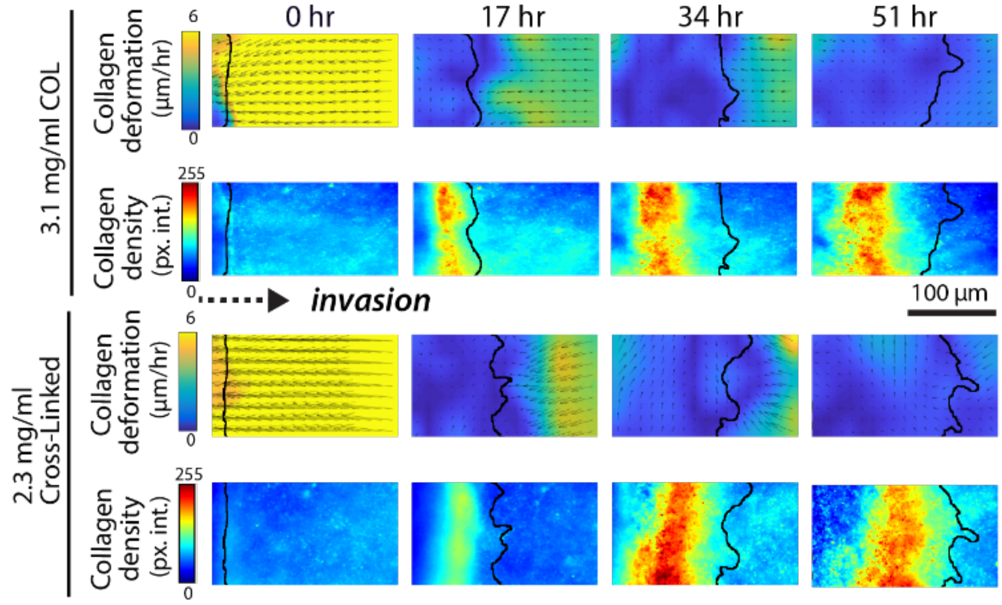


**Fig. S6. Spatial and temporal analysis of cell invasion in dense and crosslinked collagen.** Heatmaps of collagen deformation (PIV vectors) and collagen bead intensity caused by soft or stiff primed cells (black outline annotates the cell invasion front) in dense and crosslinked collagen.

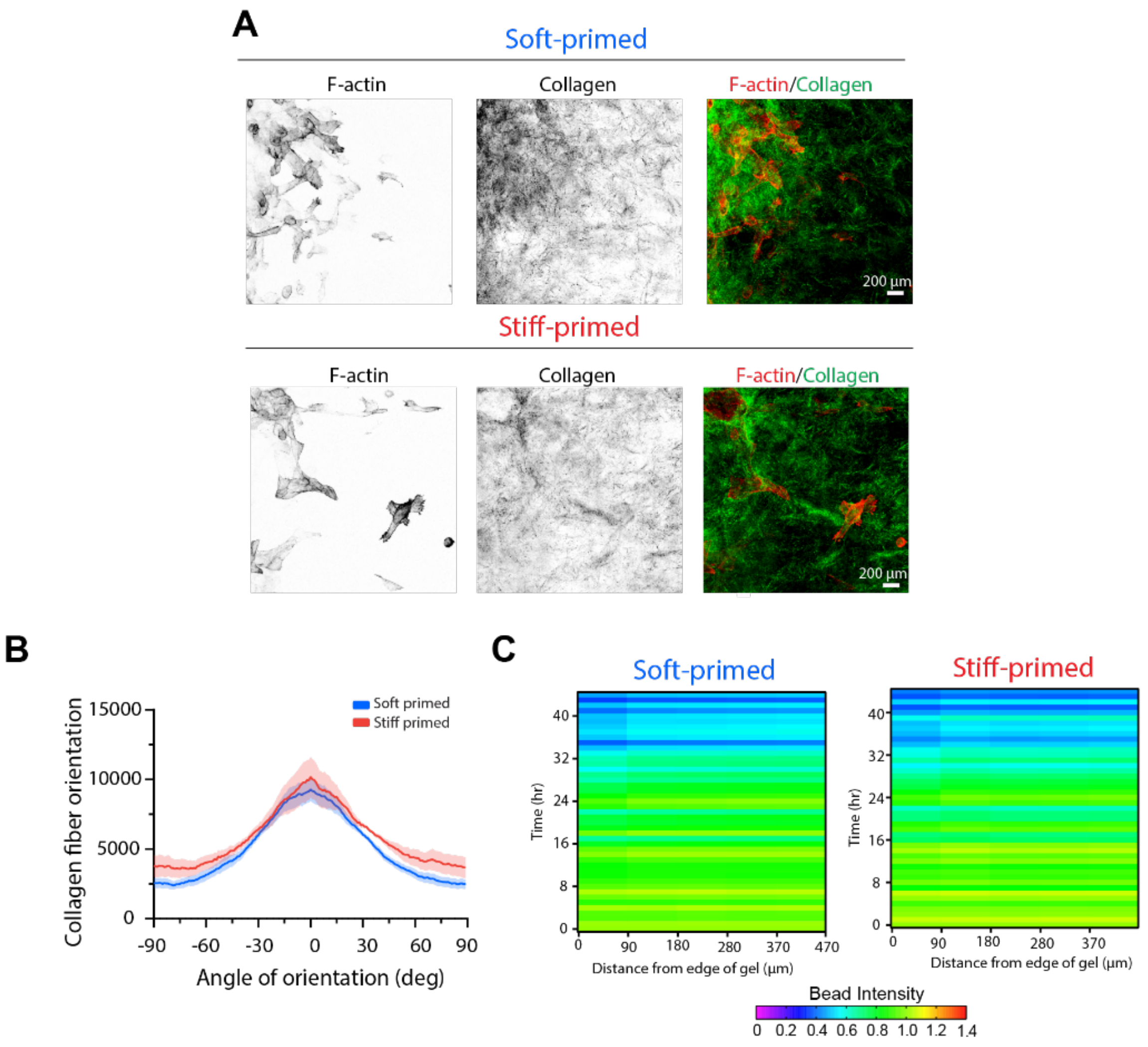


**Fig. S7. Effect of crosslinking inhibition, BAPN, on stiff and soft primed cells after 3 days of invasion. (A)** Split channel and merged images of collagen reflectance and F-actin immunofluorescence for LOX-inhibited cell invasion after priming. Scale bar, 200µm. **(B)** Alignment distribution of collagen fibers ranging from -90 to 90 degrees, where higher peaks indicate higher number of aligned fibers; showing negligible priming differences. *N=8*. **(C)** Kymographs of collagen bead intensity over time of invasion and distance from the PA gel for soft and stiff primed cells after LOX inhibition. *N=5*.

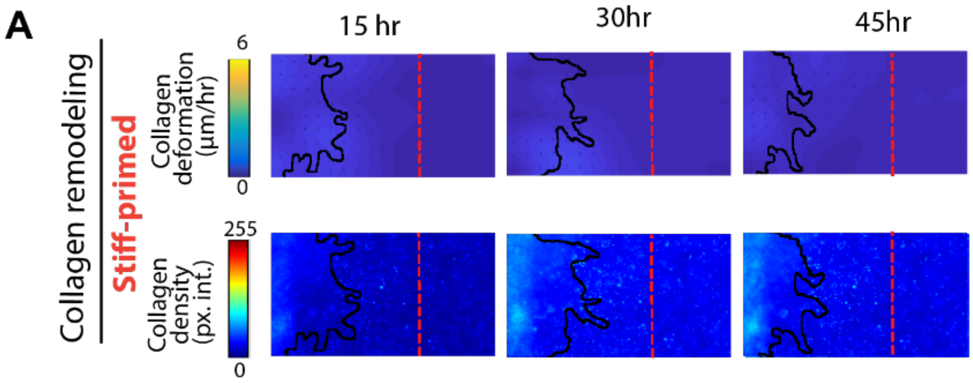


**Fig. S8. Spatial and temporal analysis of collagen remodeling after laser ablation of collagen fibers.** (**A**) Heatmaps of collagen deformation (PIV vectors) and collagen bead intensity due to laser ablation (red dotted line annotates where the collagen was cut, and black outline annotates the cell invasion front). Ablated matrix of stiff primed cells indicates reduced collagen accumulation and collagen deformation.

**Movie S1. Temporal invasion and collagen deformation of stiff-primed cells.** Stiff primed MCF10A cells expressing GFP (left panel) invading into collagen mixed with fluorescent beads (right panel). Yellow dotted lines represent the edge of the PA gel. Cells tracked for about 2 days.

**Movie S2. Temporal invasion and collagen deformation of soft-primed cells.** Soft primed MCF10A cells expressing GFP (left panel) invading into collagen mixed with fluorescent beads (right panel). Yellow dotted lines represent the edge of the PA gel. Cells tracked for about 2 days.

**Movie S3. Simulations of priming-dependent cell invasion and collagen remodeling.** Simulation showing soft-primed (left panel) and stiff-primed (right panel) epithelial cell colonies invading into collagen, overlaid with the temporally evolving spatial field of collagen remodeling $\alpha$. Simulation time is 48 hours.
